# Genetic excision of the regulatory cardiac troponin I extension in high heart rate mammal clades

**DOI:** 10.1101/2023.05.19.541292

**Authors:** William Joyce, Kai He, Mengdie Zhang, Samuel Ogunsola, Xini Wu, Kelvin T. Joseph, David Bogomolny, Wenhua Yu, Mark S. Springer, Jiuyong Xie, Anthony V. Signore, Kevin L. Campbell

**Affiliations:** Department of Biology - Zoophysiology, Aarhus University, Aarhus C, Denmark; Division of Cardiovascular Sciences, Faculty of Biology, Medicine and Health, The University of Manchester, Manchester, UK; Key Laboratory of Conservation and Application in Biodiversity of South China, School of Life Sciences, Guangzhou University, Guangzhou, Guangdong, China; Department of Physiology and Pathophysiology, University of Manitoba, Winnipeg, Manitoba, Canada; Department of Biological Sciences, University of Manitoba, Winnipeg, Manitoba, Canada; Department of Evolution, Ecology and Organismal Biology, University of California, Riverside, Riverside, United States

## Abstract

Mammalian cardiac troponin I (cTnI) contains a highly conserved N-terminal extension harboring protein kinase A targets (Ser_23/24_) which are phosphorylated during ß-adrenergic stimulation to increase cardiomyocyte relaxation rate. Here, we show that the Ser_23/24_ encoding exon 3 of *TNNI3* was pseudoexonized multiple times in shrews and moles to mimic Ser_23/24_ phosphorylation without adrenergic stimulation, facilitating the evolution of exceptionally high resting heart rates (∼1000 beats min^-1^). We further reveal alternative exon 3 splicing in distantly related bat families and that both exon 3^-^ and exon 3^+^ cTnI isoforms are incorporated into cardiac myofibrils. Finally, exon 3 of human *TNNI3* is shown to exhibit a relatively low splice strength score, offering an evolutionarily informed strategy to excise this exon to improve diastolic function during heart failure.

## Main Text

Cardiac contraction is triggered when rising cytosolic Ca^2+^ binds to troponin (Tn) in the sarcomere which, by shifting tropomyosin, allows the formation of force-generating actin-myosin cross-bridges (*1*). Thus, troponin, a trimeric complex of TnC, TnI and TnT subunits, plays a pivotal role in excitation-contraction coupling, and dysregulation or mutations in troponin subunits are associated with various cardiomyopathies and heart failure (*2*–*7*).

In mammals, the *TNNI3* gene encodes cardiac TnI (cTnI), which is differentiated from skeletal muscle troponin I paralogs (*TNNI1* and *TNNI2*) by the presence of a ∼32 amino acid N-terminal extension that plays a central regulatory role during adrenergic stimulation (*8*–*13*). In the unphosphorylated state, the N-terminal extension of cTnI interacts with cTnC to increase myofilament Ca^2+^ sensitivity (*11*). However, the N-terminal extension harbors two serine residues (Ser_23/24_) that are protein kinase A (PKA) substrates. Increased adrenergic stimulation during exercise or stress (*2, 14*) induces PKA phosphorylation of Ser_23/24_, thereby abolishing the TnI-TnC protein-protein interaction (*11, 15*) and decreasing myofilament Ca^2+^ affinity (*16*).

Accordingly, this process accelerates dissociation of Ca^2+^ during diastole and increases the rate of cardiomyocyte relaxation (*13, 17, 18*), which is essential for preserving adequate cardiac filling at elevated heart rates concomitantly driven by adrenergic stimulation (*13, 19, 20*). Deletion of the cTnI N-terminal extension in transgenic mice has been shown to mimic the effect of PKA phosphorylation (*21*–*23*) and benefit diastolic cardiac performance, particularly during aging (*10*) and deficient ß-adrenergic signaling (*24*). N-terminal cleavage of cTnI has accordingly been proposed as a potential target for the treatment of heart failure (*10, 22, 24*–*26*), when cTnI phosphorylation is often impaired due to ß_1_-adrenergic receptor desensitization (*5, 6, 27*), or in cases of familial cardiomyopathy associated with Ca^2+^ hypersensitivity (*28*).

The coding sequence of the cTnI N-terminal extension is thought to be highly conserved among eutherian mammals (*12, 29*). However, we predicted modifications of this region could reveal evolutionary adaptation in clades with exceptionally high heart rates, such as bats (order Chiroptera) and the order Eulipotyphla, which houses shrews (family Soricidae), the lineage with the highest known heart rates amongst mammals (up to 1500 beats min^-1^) (*30*–*35*).

### Genetic excision of the cTnI N-terminal extension evolved independently in shrew and mole lineages

We first collected and manually annotated *TNNI3* coding sequences from 11 eulipotyphlan (four shrews, one hedgehog, five moles, one solenodon) and six chiropteran species using NCBI GenBank (see full sequences in data S1). These data provided initial evidence for loss/inactivation of the regulatory N-terminal region (including of Ser_23/24_) encoded by exon 3 (representing an 87 bp in-frame deletion) in shrews and four of the five mole species (family Talpidae), but not bats. Linear genomic sequence comparisons illustrate that while the intronic region upstream of exon 3 is intact in shrews (fig. S1A), the sequence corresponding to exon 3 is unrecognizable, providing strong evidence this exon was inactivated in the common ancestor of Soricidae. By contrast, the sequence of this exon is identifiable in moles (fig. S1B), though exhibited splice site mutations and/or frameshift indels in all but one species initially examined. The single exception was exon 3 of the Pyrenean desman (*Galemys pyrenaicus*), a species nested within the mole clade (*36, 37*), which lacks frameshift indels, premature stop codons, and/or splice site mutations and retains an intact coding sequence with the presence of Ser_23/24_. To extend taxon sampling and verify exon 3 inactivation in Eulipotyphla, we next assembled *TNNI3* mRNA sequences from 24 eulipotyphlan species via direct cDNA sequencing (*n*=5), *de novo* transcriptome assembly (*n*=13), and/or mining of publicly available (*n*=6) cardiac transcriptomes (data S1). These data together confirm that exon 3, encoding the majority of the characteristic N-terminal extension, is not expressed in any of the shrews and moles examined, including, unexpectedly, the Pyrenean desman.

To confirm the expression of the lower molecular weight protein as predicted from genomic and mRNA sequences, western blots were initially performed with cardiac extracts from two shrews (greater white-toothed shrew, *Crocidura russula*; Eurasian pygmy shrew, *Sorex minutus*), two moles (European mole, *Talpa europaea*; Pyrenean desman, *G. pyrenaicus*), the European hedgehog (*Erinaceus europaeus*), and mouse (*Mus musculus*). We employed a TnI antibody that targets a region of the protein outside of the N-terminal extension that has previously been used to detect diverse TnI isoforms from a broad range of vertebrate species (*8*). The results confirmed the TnI protein expressed in shrew and mole/desman hearts was of a lower molecular weight (∼22 kDa) than in mouse and hedgehog (∼29 kDa) (Fig. 1B and fig. S2A), consistent with the loss of *TNNI3* exon 3 expression in the former lineages. *E. coli* expressed *G. pyrenaicus* cTnI proteins either containing or lacking the 29 amino-acid N-terminal region encoded by exon 3 further verified that interspecific differences in cTnI molecular weight can be attributed to the presence or absence of exon 3 (Fig. 1C). The European mole protein exhibited a marginally higher molecular mass than the shrew and desman proteins (Fig. 1B), attributable to the four amino acid extension at the C-terminus (data S1). Finally, to confirm that the observed bands correspond to cTnI as opposed to other TnI isoforms, in particular slow skeletal muscle troponin (ssTnI) which is expressed in fetal cardiac tissue (*38*), we next conducted mass spectrometry sequencing (LC-MS/MS) on excised ∼22 kDa bands from immunoprecipitated heart protein samples of *G. pyrenaicus* and the northern short-tailed shrew (*Blarina brevicauda*). This experiment revealed peptide matches with the predicted *TNNI3*-derived cTnI protein of both species (fig. S3), but not ssTnI. These results concur with our analysis of publicly available heart transcriptomes from two shrew (*Sorex fumeus, Blarina brevicauda*) and two mole (*Condylura cristata, Scalopus aquaticus*) species which suggested negligible (0.01-3.35%) *TNNI1* gene expression relative to *TNNI3* (table S1).

**Figure 1.**
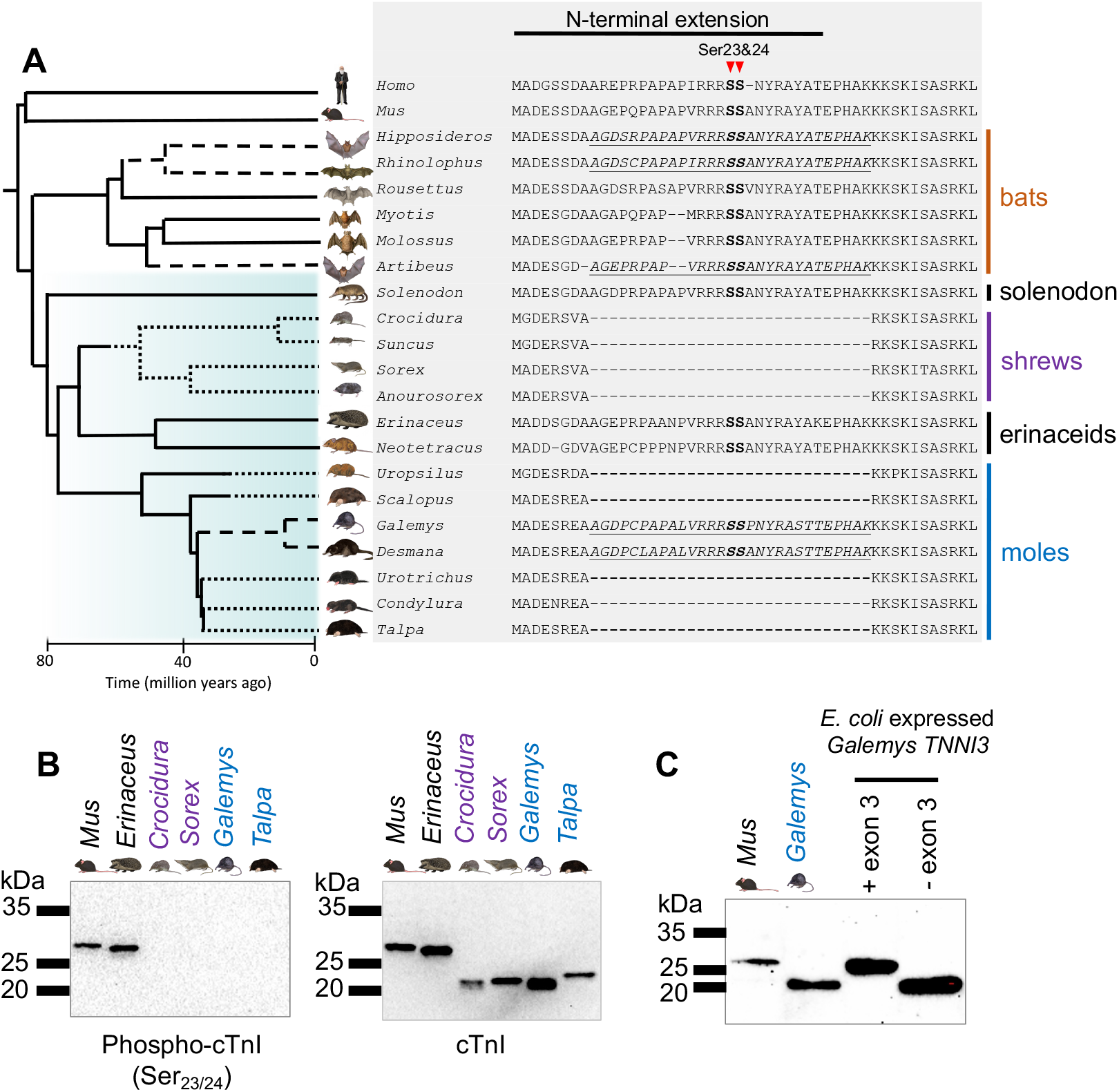
Cardiac troponin I (cTnI) primary structure, protein expression, and phosphorylation in eulipotyphlans and other mammals. (A) Representative mammalian N-terminal extension protein sequences determined from genome annotations and mRNA sequencing demonstrating the independent genomic excision of exon 3 in shrews and most mole species. Dotted lines on the cladogram show branches reconstructed to have a pseudoexonized exon 3 with the deleted exon denoted by dashes in the alignment. Dashed lines on the cladogram and underlined sequences in italics denote species whereby exon 3 is intact and supported to exhibit alternative splicing (see text for details). Protein kinase A (PKA) target residues (Ser_23/24_), which lower Ca^2+^ sensitivity of the troponin complex upon phosphorylation, are indicated in bold. *Homo*: *H. sapiens*, human; *Mus*: *M. musculus*, house mouse; *Hipposideros: H. armiger*, great roundleaf bat; *Rhinolophus*: *Rh. clivosus*, Geoffroy’s horseshoe bat; *Rousettus*: *Ro. aegyptiacus*, Egyptian fruit bat; *Myotis*: *M. myotis*, Greater mouse-eared bat, *Molossus*: *M. molossus*, Pallas’s mastiff bat; *Artibeus*: *A. jamaicensis*, Jamaican fruit bat; *Solenodon: S. paradoxus*, Hispaniolan solenodon; *Crocidura*: *C. indochinensis*, Indochinese shrew; *Suncus*: *S. etruscus*, Etruscan shrew; *Sorex*: *S. araneus*, common shrew; *Anourosorex*: *A. squamipes*, Chinese mole shrew; *Erinaceus*: *E. europaeus*, European hedgehog; *Neotetracus*: *N. sinensis*, shrew gymnure; *Uropsilus*: *U. nivatus*, snow mountain shrew-like mole; *Scalopus*: *S. aquaticus*, eastern mole; *Galemys*: *G. pyrenaicus*, Pyrenean desman; *Desmana*: *D. moschata*, Russian desman; *Urotrichus*: *U. talpoides*, Japanese shrew mole; *Condylura*: *C. cristata*, star-nosed mole; *Talpa*: *T. europaea*, European mole. (B) Western blotting of protein kinase A treated cardiac extracts from mouse and five eulipotyphlan species. The membrane was first probed for Ser_23/24_ phoshorylated-cTnI, then stripped and re-probed for total cTnI. *Mus*: *M. musculus*, house mouse; *Erinaceus*: *E. europaeus*, European hedgehog; *Crocidura*: *C. russula*, greater white-toothed shrew; *Sorex*: *S. minutus*, pygmy shrew; *Galemys*: *G. pyrenaicus*, Pyrenean desman; *Talpa*: *T. europaea*, European mole. For blots incorporating additional sample replicates without PKA treatment and using direct cTnI blotting, see fig. S1. (C) Western blot of *Escherichia coli* expressed *G. pyrenaicus TNNI3* containing and lacking exon 3 sequence, alongside *M. musculus* and *G. pyrenaicus* heart homogenates, verifying that the observed interspecific differences in cTnI molecular weights can be attributed to the presence or absence of exon 3. Image credits: Carl Buell: *Homo, Solenodon, Sorex, Erinaceus*; Laura Cadiz: *Mus, Hipposideros, Artibeus, Crocidura, Suncus, Scalopus, Galemys, Talpa*; Umi Matsushita: *Anourosorex, Neotetracus, Uropsilus, Desmana, Urotrichus, Condylura*. Wikimedia Commons (public domain): *Rhinolophus, Rousettus, Myotis, Molossus*.

We additionally treated protein extracts from the initial six species with PKA (recombinant catalytic subunit) *in vitro* and performed western blots to detect Ser_23/24_ phosphorylated cTnI. As expected, only the hedgehog and mouse exhibited bands that were co-localized with cTnI (Fig. 1B), reaffirming that the cTnI protein of shrews and moles lacks the N-terminal extension PKA phosphorylation sites. We subsequently performed western blots using an antibody that detects all phosphorylated PKA substrates (“RRXS*/T*”) on the same PKA-treated extracts. This antibody also detected strong bands in hedgehog and mouse at ∼29 kDa, consistent with cTnI, but did not reveal PKA phosphorylated proteins in the shrew or mole species at the expected location of cTnI (fig. S2B). The general PKA substrate antibody did, however, detect a range of other (unidentified) PKA substrates in moles and shrews (fig. S2B), indicating that these species have undergone a selective loss of cTnI phosphorylatability, as opposed to showing a generalized lack of responsiveness to PKA.

In contrast to shrews and moles, genomic, mRNA, and/or protein evidence together support the presence of the cTnI N-terminal extension PKA phosphorylation residues in all four erinaceid (hedgehog and gymnure) species investigated (*Atelerix albiventris, E. europaeus, Neohylomys hainanensis*, and *Neotetracus sinensis*) (data S1). As phylogenomic data indicate a robust sister relationship between erinaceids and shrews (*36, 37*), our results strongly support the independent genetic excision of *TNNI3* exon 3 in the shrew and mole lineages (Fig. 1A). The deletion of *TNNI3* exon 3 is analogous to the N-terminal cleavage previously characterized in transgenic mice (*10, 22*) in that they both permanently mimic the effects of PKA-mediated Ser_23/24_ phosphorylation. Thus, this evolutionary innovation likely contributes to improving diastolic performance by augmenting ventricular filling time and hence facilitated the evolution of exceptionally high—up to 1000 beats min^-1^ (*30, 31, 39*)—resting heart rates in shrews. Specifically, we posit that the evolution of high mass-specific metabolic rates in early (Eocene) members of this lineage were supported by progressive increases in resting heart rate that eventually favored exon 3 skipping to enhance diastolic function. Thereafter relaxed selection on this exon presumably led it to become permanently inactivated, fostering the evolution of even higher resting heart rates. It was initially surprising to find a genetically abbreviated N-terminal extension in mole hearts, given that fossorial European (*T. europaea*) and eastern moles (*Scalopus aquaticus*) have relatively slow heart rates for their body mass (∼160 and ∼230 beats min^-1^, respectively (*40, 41*)). However, strictly fossorial Talpini and Scalopini moles evolved convergently from small, ambulatory, and presumably highly active shrew-like ancestors (*36, 37*). Indeed, American shrew moles (*Neurotrichus gibbsii*) and star-nosed moles (*Condylura cristata*), which are nested between the above two fossorial clades (*36*), exhibit high mass-specific metabolic rates that are comparable to soricid shrews (*42, 43*). Thus, it is plausible that exon 3 of *TNNI3* was independently pseudoexonized in small, shrew-like mole ancestors of fossorial Talpini and Scalopini under much the same selective pressures as in shrews, i.e., to support extreme cardiac and metabolic demands.

To better elucidate the evolutionary and genomic underpinnings of exon 3 inactivation within the talpid family, we sequenced the *TNNI3* N-terminal region (using genomic DNA) from three additional mole species (*Uropsilus gracilis, Urotrichus talpoides*, and *Desmana moschata*). These data reveal that, like its close relative *G. pyrenaicus*, the Russian desman (*D. moschata*) retains an intact exon 3 that is potentially functional/expressed (fig. S4). By contrast, representative species from the remaining five talpid clades (*Talpa, Condylura, Urotrichus, Scalopus, Uropsilus*) all exhibit inactivating mutations in exon 3 that are hallmarks of pseudoexonization (fig. S4). For example, the exon 3 donor and acceptor splice sites are intact in Scalopini moles and Urotrichini shrew moles, though their exons have been functionally inactivated by frameshift insertions/deletions. *Uropsilus* and *Condylura*, in turn, have independently accumulated acceptor splice site mutations and frameshift deletions. Finally, Talpini moles exhibit a 10-bp coding sequence frameshift deletion and a 26-bp deletion that encompasses the acceptor splice site. The central placement of desmans within Talpidae (*36*) implies the loss of regulatory cTnI phosphorylation must have occurred a minimum of three times in the common ancestors of: (1) shrew-like moles (*Uropsilus*), (2) Scalopini moles (*Scalopus*), and (3) a clade that includes Talpini moles (*Talpa*), star-nosed moles (*Condylura*), and shrew moles (*Urotrichus*). However, as none of the above talpid lineages share common inactivating mutations of exon 3 (fig. S4), the N-terminal extension of cTnI appears to have been independently disrupted in *Uropsilus, Scalopus, Talpa, Condylura*, and *Urotrichus* (see cladogram of Fig. 1A).

### Desman *TNNI3* exon 3 evolved under purifying selection and may be alternatively spliced

Given our finding that exon 3 is intact in desmans but inactivated in shrews and other moles, we performed selection analyses with codeml (*35*) and RELAX (*55*) to probe the molecular evolutionary history of *TNNI3* in these eulipotyphlans versus other placental mammals. First, we examined complete protein-coding sequences, minus exon 3, for a data set comprised of 48 taxa including 16 shrews and eight moles (fig. S5). This allowed us to test the hypothesis that exons 1, 2, and 4-8 have remained under purifying selection in shrew and mole taxa with pseudoexonized exon 3 sequences. The results of these analyses confirm that the *TNNI3* gene

(with exon 3 removed) has evolved under strong purifying selection within Eutheria including moles and shrews (ω=0.038 to 0.067 with different codon frequency models) (fig. S5 and data S2A). Statistical tests reject the hypothesis that ω values in shrews and non-desman moles are different from those in other placental mammals (data S2A,C). Further, ω values for *TNNI3* coding sequences (minus exon 3) for different shrew and mole branches with inactivating mutations in exon 3 range from 0.0001 to 0.1846 and are fully consistent with purifying selection (data S2B). By contrast, ω values for pseudoexonic branches of exon 3 in non-desman moles are under relaxed selection (ω =0.867) relative to non-talpid mammals (ω=0.142) (p=0.000), as may be expected for pseudoexonic sequences (Fig. 2, fig. S6 and data S2D). In the case of desmans, we hypothesized that the coding sequences for exon 3 may be evolving under neutral selection— like in other talpid species—given the absence of direct evidence for exon 3 expression in mature mRNA or heart western blots (Fig. 1B and fig. S2C). To test this hypothesis, we compared ω values in Desmanini to non-talpid placental mammals. The ω value for desmans (0.490) is elevated, but not significantly (p=0.07) different from the ω value in other mammals (0.121) with an intact copy of exon 3 (fig. S7 and data S2E). It is noteworthy in this regard that the desman exon 3 still encodes the highly conserved PKA target motif (“RRRS”; Fig. 1A and data S1). To validate that desman Ser_23/24_ are still potential PKA substrates, we used NetPhos (*44*) to predict phosphorylation targets in mouse, human, European hedgehog and desman *TNNI3*. This analysis showed that the PKA target scores for desman cTnI Ser_23/24_ are on a par with other mammals (fig. S8), consolidating that the *TNNI3* exon 3, including the PKA motif, has remained remarkably well conserved in this lineage.

**Figure 2.**
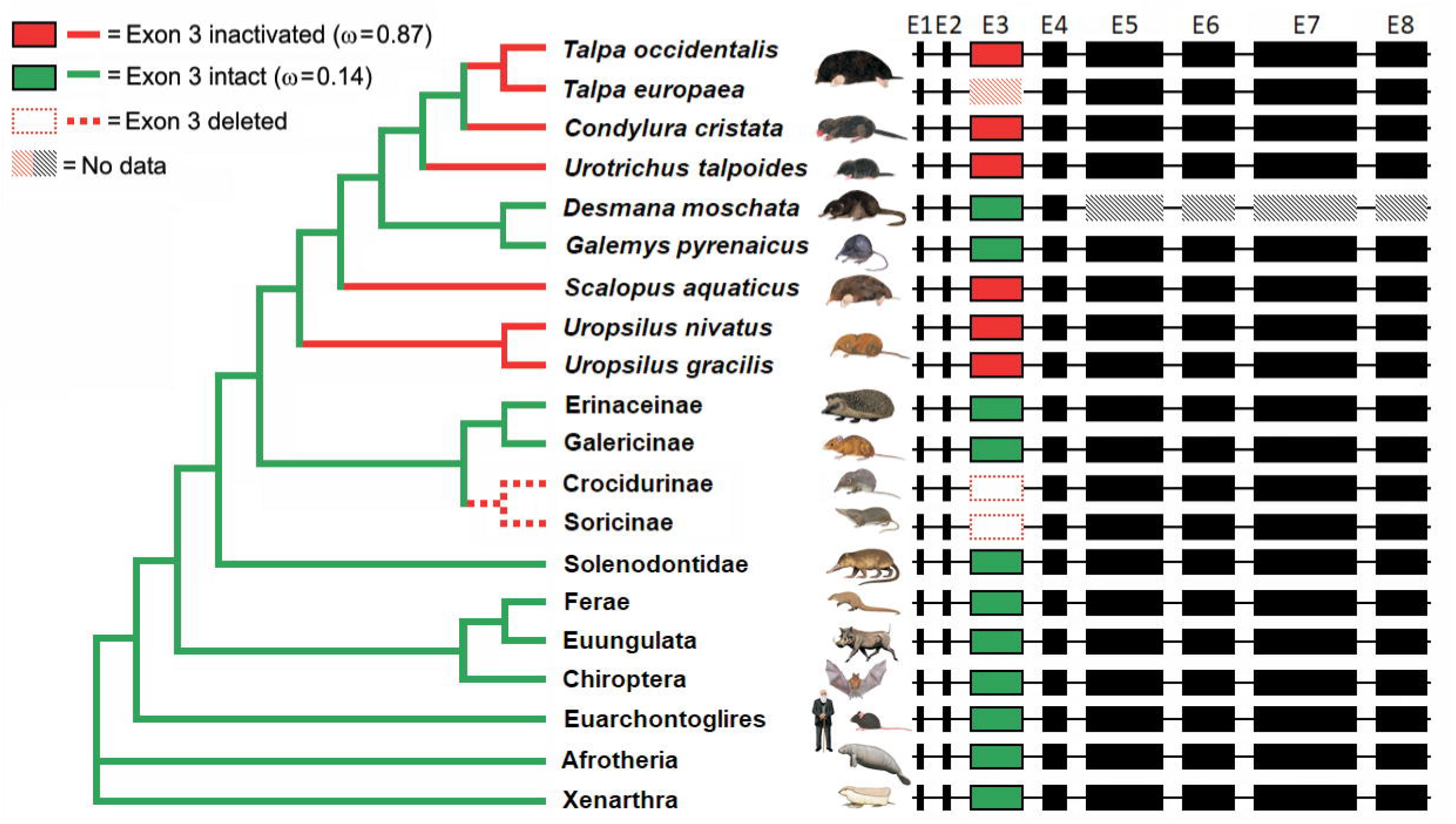
Phylogenetic tree and exon diagrams showing different evolutionary pressures acting on *TNNI3* exon 3 in a diverse array of placental mammals as evidenced by inactivating mutations and selection analyses (ω values). Green lines denote branches whereby exon 3 is intact (green boxes) and evolving under purifying selection (ω=0.14) while red lines denote branches with inactivating mutations in exon 3 (red boxes) and evolving under relaxed selection (ω=0.87, see data S2D). Exon 3 of all examined shrew species was not included in this analysis as it was unrecognizable (dashed red boxes) and was presumably inactivated in the common ancestor of Soricidae (dotted red lines). By contrast, the coding sequence of *TNNI3* exons 1, 2, 4-8 (black boxes) evolved under strong purifying selection in all placental mammals (ω=0.052) (data S2A). Hatched boxes represent (pseudo)exons where sequence data are unavailable. For complete information on the species used in the RELAX (*55*) selection analyses summarized here, refer to Figs. S5-S7. The species tree is based on Meredith et al. (2011) (*76*) and He et al. (2021) (*36*). Abreviations, E = exon. Image credits: Carl Buell: Erinaceinae, Soricinae, Solenodontidae, Ferae, Euungulata, human (Euarchontoglires), Afrotheria, Xenarthra; Laura Cadiz: *Talpa, Galemys, Scalopus*, Crocidurinae, Chiroptera, mouse (Euarchontoglires); Umi Matsushita: *Condylura, Urotrichus, Desmana, Uropsilus*, Galericinae.

Results of the selection analyses are consistent with the hypothesis that exon 3 of desmans could be variably expressed (i.e., via alternative exon splicing). To further examine the potential for *TNNI3* exon 3 alternative splicing in this clade, the strengths of the upstream (intron 2) 3’ acceptor splice site and downstream (intron 3) 5’ donor splice site were assessed using maximum entropy modelling (*45*) with comparisons to representative outgroup species with intact exon 3 sequences (Fig. 3 and data S3). The splice strengths (MaxEnt scores) for the upstream and downstream splice sites of exon 2 and exon 4 were relatively high across all species. However, only desmans exhibited drastically reduced 3’ splice site strengths for exon 3 (Fig. 3B). The desman 5’ *TNNI3* exon 3 splice site, by contrast, was highly conserved and retained a comparably high splice strength score relative to the other eulipotyphlan species (solenodons and erinaceids) (Fig. 3E). Notably, this combination of a low 3’ splice score and a high 5’ splice score renders exon 3 of desmans liable to alternative splicing (*46*). This conclusion conflicts with the prevailing dogma that *TNNI3* does not normally undergo alternative exon splicing in vertebrates (*12, 47, 48*). Although cTnI protein assays from five *G. pyrenaicus* specimens failed to substantiate alternative exon splicing in cardiac tissue (fig. S2C), it remains possible that exon 3 is expressed in the heart under certain circumstances or in other tissues. For example, the human protein atlas provides evidence for cTnI expression in testis tissue (*49*). Indeed, publicly available RNA-seq data from *T. occidentalis* testis tissue (*50*) confirms this expression profile in Talpini moles, thus also opening the possibility for *TNNI3* exon 3 translation in this organ of desmans. Furthermore, various other non-canonical roles for troponins are emerging (*51*), for instance the expression of *TNNI3* has been reported in cancer cells (*51*–*53*) and rat brain endothelial cells (*54*) as well as in the nucleus of striated muscle (*55*), and our findings contribute to the growing understanding of the complexity of *TNNI3* expression. However, it is also possible that *TNNI3* exon 3 skipping was genetically assimilated either independently or in a common ancestor of desmans and that inactivating mutations in the exon have not yet become fixed.

**Figure 3.**
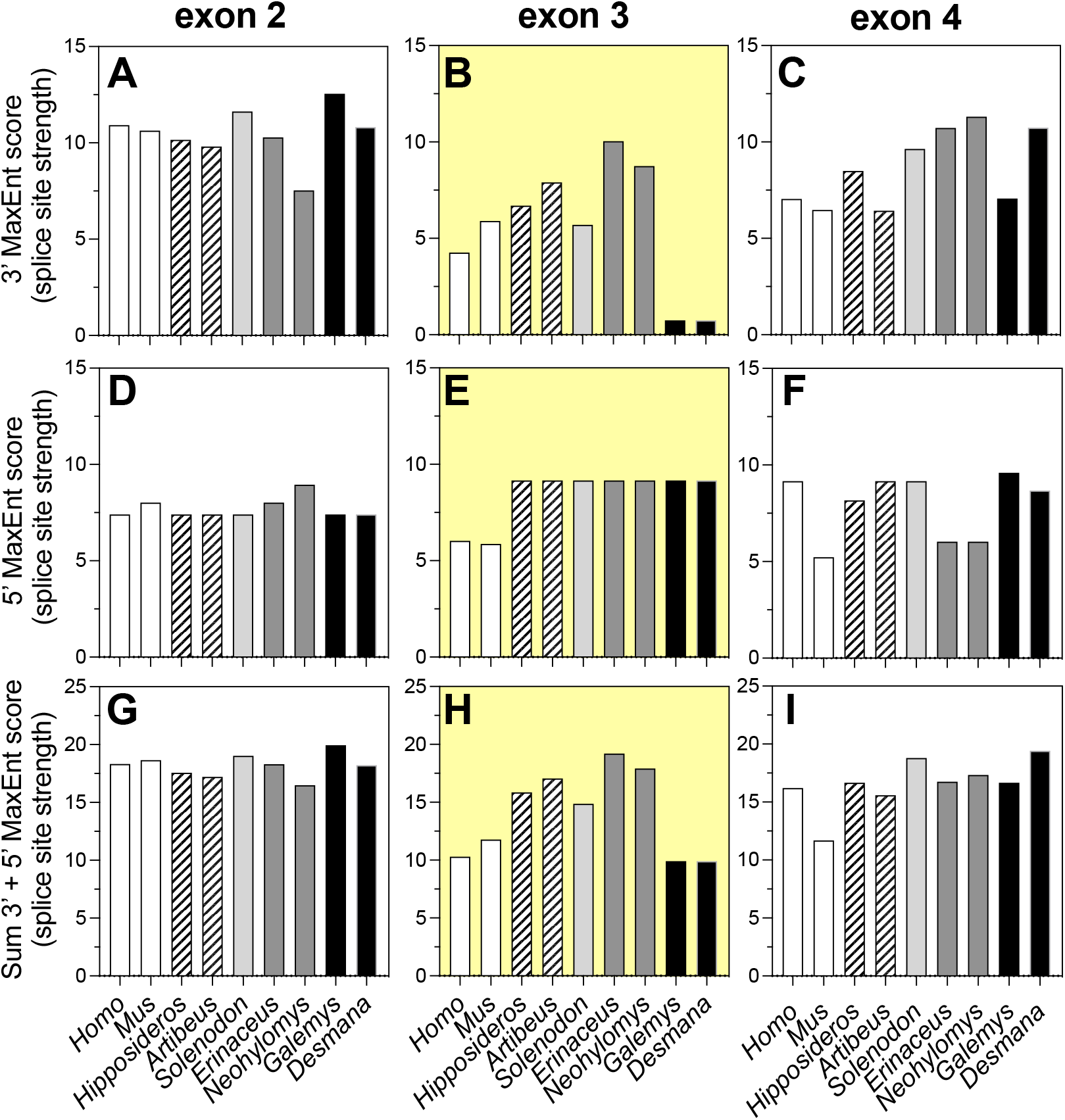
Maximum entropy (MaxEnt) upstream (3’ site of previous intron) and downstream (5’ site of subsequent intron) splice site strength scoring for *TNNI3* (A) exon 2, (B) exon 3, and (C) exon 4. This analysis was conducted for desmans (*Galemys*: *G. pyrenaicus*, Pyrenean desman; *Desmana: D. moschata*, Russian desman), the only talpid moles with an intact *TNNI3* exon 3, and outgroups that also express exon 3 (*Homo*: *H. sapiens*, human; *M. musculus*, house mouse; *Hipposideros: H. armiger*, great roundleaf bat; *Artibeus*: *A. jamaicensis*, Jamaican fruit bat; *Solenodon: S. paradoxus*, Hispaniolan solenodon; *Erinaceus*: *E. europaeus*, European hedgehog; *Neohylomys*: *N. hainanensis*, Hainan gymnure). Splice site strength was well conserved and high across species for exons 2 (A, D, G) and 4 (C, F, I). However, desmans exhibited dramatically reduced 3’ splice site strength for exon 3 (B), although they retained high 5’ splice site strength (E), which together indicates the exon is likely capable of alternative splicing (*46*).

### *TNNI3* exon 3 is alternatively spliced in bats

Like shrews, chiropteran bats are able to sustain high mass-specific metabolic rates (*56*) with some species exhibiting maximal heart rates up to 1000 beats min^-1^ (*57, 58*). While our initial genomic data showed that the phosphorylatable cTnI N-terminal extension of bats is intact (data S1), subsequent analysis of publicly available cardiac *TNNI3* transcripts from 11 chiropteran species unambiguously demonstrates alternative splicing of exon 3 in multiple species from representatives of two distantly related clades (Phyllostomidae, suborder Yangochiroptera, and Hipposideridae/Rhinolophidae, suborder Yinpterochiroptera) (Fig. 1A and fig. S9). As the abundance of mRNA transcripts lacking exon 3 were estimated to comprise from 31-51% of total *TNNI3* in members of both clades (data from our assembly of publicly available heart transcriptomes; table S2), we collected two great roundleaf bat (*Hipposideros armiger*) specimens to verify translation and incorporation of the two different cTnI isoforms into sarcomeres (Fig. 4). These experiments revealed that the lower molecular weight cTnI formed 21.0 ± 6.8% (mean ± SD) of total cTnI in cardiac homogenates and 31.2 ± 2.0% of total cTnI in myofibrils (data S4), confirming the alternatively spliced cTnI isoform is well placed to contribute to cardiac function. As was the case with shrews/moles, cardiac *TNNI1* transcript levels were extremely low (<0.5%) relative to *TNNI3* in all bat species exhibiting alternative exon 3 splicing (table S1), providing strong evidence that both bands correspond to cTnI. Protein sequencing (LC-MS/MS) with parallel reaction monitoring verifies this conclusion (fig. S10A) and further confirms exon 3 skipping in the ∼22 kDa *H. armiger* band (i.e., the exon 2/exon 4 splice boundary is present). Similarly, the exon 2/exon 4 splice boundary was also evident in the ∼22 kDa Asian house shrew (*Suncus murinus*) cTnI protein analyzed with the same method (fig. S10B).

**Figure 4.**
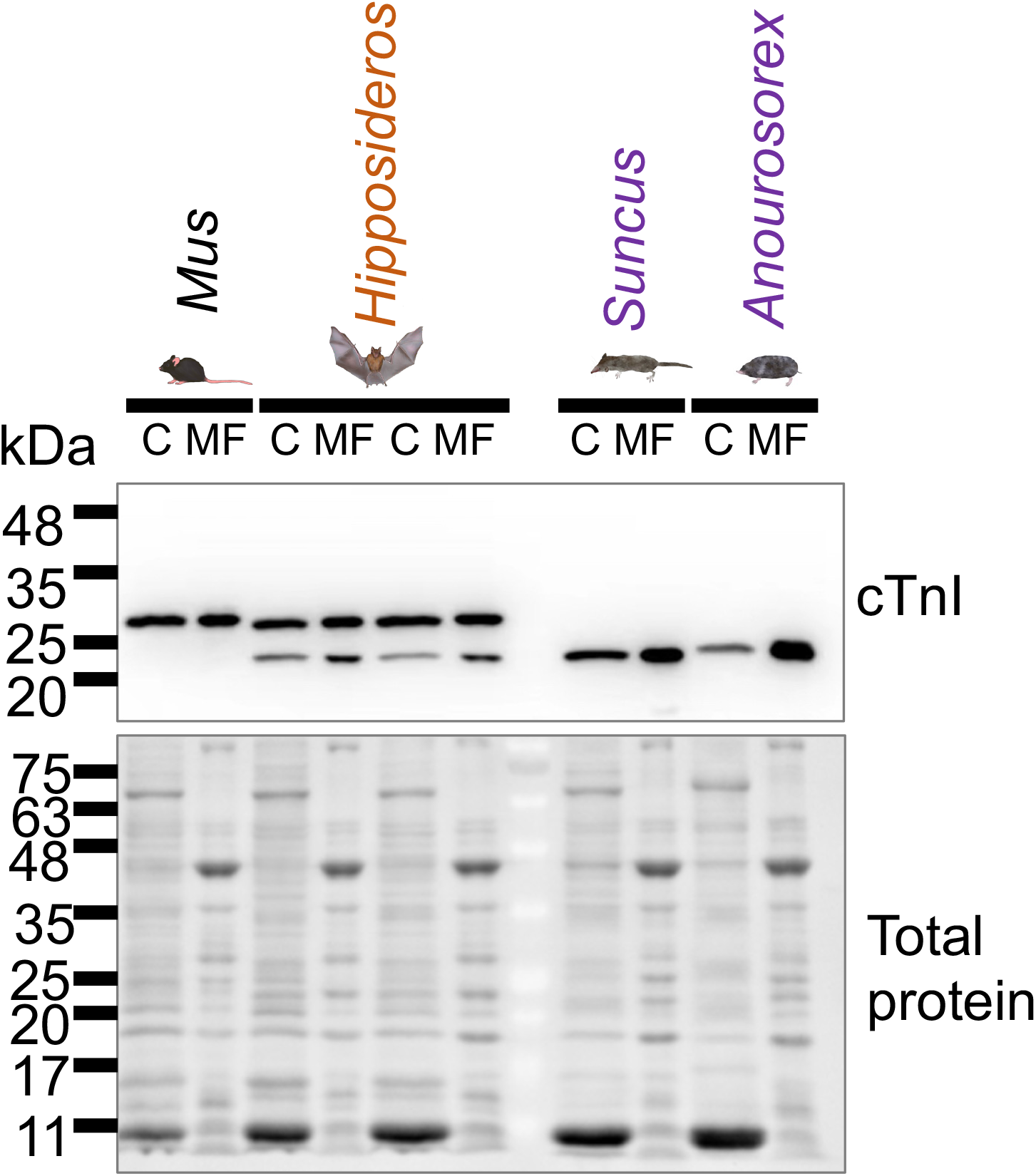
Alternatively spliced cTnI protein isoforms are expressed and incorporated into heart myofibrils of great roundleaf bats. A Western blot for troponin expression was performed on crude cardiac homogenates (C) or washed myofibril extracts (MF) of ventricles from: *Mus*: *M. musculus*, house mouse; *Hipposideros: H. armiger*, great roundleaf bat (2 individuals); *Suncus*: *S. murinus*, Asian house shrew; *Anourosorex*: *A. squamipes*, Chinese mole shrew. Note that *H. armiger* exhibits both ∼22 and ∼29 kDa bands, consistent with *TNNI3* alternative splicing, as identified in the transcriptome of this species. Notably, the lower molecular weight (∼22 kDa) cTnI protein is also demonstrated to be incorporated into the cardiac sarcomeres of soricid and crocidurian shrews. Image credits: Laura Cadiz: *Mus, Hipposideros, Suncus*; Umi Matsushita: *Anourosorex*.

The finding of alternative exon 3 splicing in bats supports our proposed scenario for the observed pattern of exon 3 pseudoexonization in shrews and moles. Specifically, the ability to alternatively splice exon 3 may have had ancient evolutionary origins in the common ancestors of these latter clades to support evolutionary increases in heart rate. The presence of intact exon 3 donor and acceptor splice sites in several extant mole lineages (i.e., *Scalopus* and *Urotrichus*) provides evidence for the genetic assimilation of exon 3 skipping in pre-mRNA transcripts, which was followed by the independent degeneration of this coding region in each of the five non-desman mole clades (Fig. 1 and fig. S4). Surprisingly, exon 3 of two representative bat species shown to be capable of alternatively splicing, *Hipposideros armiger* and *Artibeus jamaicensis* (fig. S9), exhibits high 3’ and 5’ MaxEnt splice site scores (Fig. 3), in favor of only exon inclusion (*46*). This suggests that exon 3 alternative splicing must be mainly dependent on *trans*-acting factors (*59*) in these species.

We hypothesize that the genomic retention of *TNNI3* exon 3 in Chiroptera may in part be a preadaptation that facilitates the ability of many bat species to enter into seasonal hibernation and/or daily torpor, which, when contrasted with the energetic demands of powered flight, results in an extraordinary heart rate scope (*60, 61*). Low body temperature, as experienced during hibernation, reduces myofilament Ca^2+^ sensitivity (*62, 63*), potentially leading to cardiac asystole. However, in compensation, some hibernating mammals are able to increase myofilament Ca^2+^ affinity (*64*). This is likely attributable, at least in part, to the effects of reduced adrenergic stimulation (*65*) and a corresponding reduction of cTnI phosphorylation. In this situation, the non-phosphorylated N-terminal extension would play an essential role to safeguard myofilament Ca^2+^ sensitivity. Although the ability to enter short-term torpor is widespread among crocidurine (white-toothed) shrews (*66*), no other eulipotyphlan lineage apart from hedgehogs (which express the phosphorylatable N-terminal extension) is known to exploit true hibernation. This observation suggests that the loss of *TNNI3* exon 3 could be a historical contingency that contributes to the inability of moles and shrews to partake in hibernation as a seasonal energy conservation strategy.

### Tachycardic mammals reveal evolutionarily informed pathways to therapeutically target cTnI

Reduced phosphorylation of the cTnI N-terminal extension is a hallmark of chronic heart failure that increases Ca^2+^ affinity of the troponin complex and may contribute to diastolic cardiac dysfunction (*5, 6, 27*). However, the transgenic overexpression of N-terminal truncated cTnI in mice has been shown to mimic the effect of PKA activation in reducing myofilament Ca^2+^ sensitivity (*10, 22*). As such, removal of the N-terminal extension of cTnI is already considered a potential treatment to restore diastolic function during heart failure (*10, 22, 24, 25*). Crucially, in the absence of chronic adrenergic stimulation, which is known to lead to ß_1_-adrenergic receptor desensitization and cardiac hypertrophy during the progression of heart failure (*67*), N-terminal truncation provides a mechanism to increase the rate of myofibril relaxation and improved diastolic filling of the heart (*22*).

Although generated by a different mechanism—pseudoexonization of exon 3 in shrews and moles *versus* insertion of an artificial translation initiation codon in transgenic mice (*22*)— the shortened N-terminal cTnIs share a close resemblance (fig. S11). The natural capacity for alternative splicing of exon 3, here evidenced in bats and implicated in desmans, thus uncovers a reversible strategy to induce exon 3 exclusion in mature human *TNNI3* transcripts. Although the *TNNI3* exon 3 was not known to be alternatively spliced in any mammal prior to this study (*12, 48*), and we can find no evidence that it is ever normally excluded in human *TNNI3*, our maximum entropy modelling indicates exon 3 has relatively low splice site strength scores (Fig. 3B,E,H). Indeed, the sum MaxEnt score of human *TNNI3* exon 3 (i.e., 3’ + 5’ MaxEnt score: 10.29) is much lower than the two neighboring exons and only slightly exceeds that of desmans (9.91), which we have demonstrated ordinarily skip exon 3. This combination of 3’ and 5’ splice site scores of human *TNNI3* exon 3 is predicted to result in only a ∼66% likelihood of exon inclusion (*46*). Moreover, we have found a common (gnomAD frequency: 35.1%; TOPMed frequency: 37.8%) single nucleotide polymorphism (NCBI SNP: rs3729836) at the upstream 3’ splice site of human *TNNI3* exon 3 (data S3), which lowers the 3’ MaxEnt score even further (from 4.26 to 3.69) and reduces the likelihood of exon 3 inclusion to ∼57%. This SNP also specifically disrupts the human branch point consensus sequence (YTNAY; (*68*)) in pre-mRNA so may be particularly predisposed to the induction of alternative splicing (*69*). Together, these results suggest that although *TNNI3* exon 3 alternative splicing is not naturally exploited in our own species, it carries genomic signatures that may render it amenable to the induction of exon 3 skipping, for example *via* genetic strategies such as antisense oligonucleotides (*70, 71*) or with small molecule approaches (*72*). These strategies are preferable to (post-translational) proteolytic cleavage of the N-terminus, the context of extensive previous work (*10, 21, 22, 24*–*26*), given the high rate of sarcomeric protein turnover in heart muscle and low specificity of proteases—such as calpains (*26, 73*)—capable of cleaving cTnI (*74*).

By uncovering exon 3 alternative splicing in diverse species such as bats and potentially desmans, our study lays the foundations to explore how exon 3 excision might be therapeutically induced during diastolic heart failure in humans. Although additional preclinical research is warranted, this approach may be relevant to treating inherited cardiomyopathies associated with increased myofilament Ca^2+^ sensitivity such as hypertrophic cardiomyopathy though would not be appropriate in conditions associated with decreased myofilament Ca^2+^ sensitivity for example dilated cardiomyopathy (*2*) or hypertrophic cardiomyopathy associated with mutations at the exon-intron border or resulting in a reading frameshift (*75*). In sum, by studying the hearts of small mammals with exceptionally high heart rates (*30*), we have shown that nature realized the solution to habitually accelerate diastolic relaxation long before modern biomedicine.

## Acknowledgments

We are grateful to Jorge Gonzales Esteban for providing Pyrenean desman heart samples and five anonymous donors for additional tissue samples used in this study. We thank Yinxu Wang for helping with transcriptome assembly and Xiongjun Wang for assistance with the western blot and Co-IP experiments. Animal illustrations were created by Laura Cadiz, Umi Matsushita, and Carl Buell (authorization for the latter kindly provided by John Gatesy) and used with permission.

## Author contributions

Conceptualization: WJ, KLC

Methodology: WJ, KH, MZ, SO, XW, KTJ, DB, JX, MSS, AVS, KLC

Investigation: WJ, KH, MZ, SO, XW, KTJ, DB, WY, JX, MSS, AVS, KLC

Visualization: WJ, KH, MSS, KLC

Funding acquisition: WJ, KH, JX, KLC

Project administration: WJ, KLC

Supervision: KLC, KH, JX

Writing – original draft: WJ, KLC

Writing – review & editing: WJ, KH, DB, JX, MSS, AVS, KLC

## Supplementary Materials

Materials and Methods (including Tables S3 and S4)

Tables S1 and S2

Figs. S1 to S10

References (*74-83)*

## References

1. D. M. Bers, Cardiac excitation–contraction coupling. Nature 415, 198–205 (2002).

2. J. van der Velden, G. J. M. Stienen, Cardiac disorders and pathophysiology of sarcomeric proteins. Physiol. Rev. 99, 381–426 (2019).

3. S. Morimoto, Sarcomeric proteins and inherited cardiomyopathies. Cardiovasc. Res. 77, 659–666 (2008).

4. P. M. Hwang, B. D. Sykes, Targeting the sarcomere to correct muscle function. Nat. Rev. Drug Discov. 14, 313–328 (2015).

5. P. J. M. Wijnker, A. M. Murphy, G. J. M. Stienen, J. van der Velden, Troponin I phosphorylation in human myocardium in health and disease. Neth. Heart J. 22, 463–469 (2014).

6. G. S. Bodor, A. E. Oakeley, P. D. Allen, D. L. Crimmins, J. H. Ladenson, P. A. Anderson, Troponin I phosphorylation in the normal and failing adult human heart. Circulation 96, 1495–1500 (1997).

7. H. J. Tadros, C. S. Life, G. Garcia, E. Pirozzi, E. G. Jones, S. Datta, M. S. Parvatiyar, P. B. Chase, H. D. Allen, J. J. Kim, J. R. Pinto, A. P. Landstrom,Meta-analysis of cardiomyopathy-associated variants in troponin genes identifies loci and intragenic hot spots that are associated with worse clinical outcomes. J. Mol. Cell. Cardiol. 142, 118–125 (2020).

8. W. Joyce, D. M. Ripley, T. Gillis, A. C. Black, H. A. Shiels, F. G. Hoffmann, A revised perspective on the evolution of troponin I and troponin T gene families in vertebrates. Genome Biol. Evol., evac173 (2022).

9. H. E. Salhi, V. Shettigar, L. Salyer, S. Sturgill, E. A. Brundage, J. Robinett, Z. Xu, E. Abay, J. Lowe, P. M. L. Janssen, J. A. Rafael-Fortney, N. Weisleder, M. T. Ziolo, B. J. Biesiadecki, The lack of Troponin I Ser-23/24 phosphorylation is detrimental to in vivo cardiac function and exacerbates cardiac disease. J. Mol. Cell. Cardiol., doi: 10.1016/j.yjmcc.2023.01.010 (2023).

10. B. J. Biesiadecki, K. Tachampa, C. Yuan, J.-P. Jin, P. P. de Tombe, R. J. Solaro, Removal of the cardiac troponin I N-terminal extension improves cardiac function in aged mice. J. Biol. Chem. 285, 19688–19698 (2010).

11. P. M. Hwang, F. Cai, S. E. Pineda-Sanabria, D. C. Corson, B. D. Sykes, The cardiac-specific N-terminal region of troponin I positions the regulatory domain of troponin C. Proc. Natl. Acad. Sci. U. S. A. 111, 14412–14417 (2014).

12. J.-J. Sheng, J.-P. Jin, TNNI1, TNNI2 and TNNI3: Evolution, regulation, and protein structure–function relationships. Gene 576, 385–394 (2016).

13. S. Marston, Recent studies of the molecular mechanism of lusitropy due to phosphorylation of cardiac troponin I by protein kinase A. J. Muscle Res. Cell Motil., doi: 10.1007/s10974-022-09630-4 (2022).

14. D. M. Bers, Y. K. Xiang, M. Zaccolo, Whole-Cell cAMP and PKA activity are epiphenomena, nanodomain signaling matters. Physiol. Bethesda Md 34, 240–249 (2019).

15. G. Ferrières, M. Pugnière, J. C. Mani, S. Villard, M. Laprade, P. Doutre, B. Pau, C. Granier, Systematic mapping of regions of human cardiac troponin I involved in binding to cardiac troponin C: N- and C-terminal low affinity contributing regions. FEBS Lett. 479, 99–105 (2000).

16. J. Wattanapermpool, X. Guo, R. J. Solaro, The unique amino-terminal peptide of cardiac troponin I regulates myofibrillar activity only when it is phosphorylated. J. Mol. Cell. Cardiol. 27, 1383–1391 (1995).

17. S. Yasuda, P. Coutu, S. Sadayappan, J. Robbins, J. M. Metzger, Cardiac transgenic and gene transfer strategies converge to support an important role for troponin I in regulating relaxation in cardiac myocytes. Circ. Res. 101, 377–386 (2007).

18. R. Zhang, J. Zhao, A. Mandveno, J. D. Potter, Cardiac troponin I phosphorylation increases the rate of cardiac muscle relaxation. Circ. Res. 76, 1028–1035 (1995).

19. W. Joyce, T. Wang, How cardiac output is regulated: August Krogh’s proto-Guytonian understanding of the importance of venous return. Comp. Biochem. Physiol. A. Mol. Integr. Physiol. 253, 110861 (2021).

20. S. Marston, J. R. Pinto, Suppression of lusitropy as a disease mechanism in cardiomyopathies. Front. Cardiovasc. Med. 9 (2023).

21. L. K. Gunther, H.-Z. Feng, H. Wei, J. Raupp, J.-P. Jin, T. Sakamoto, Effect of N-terminal extension of cardiac troponin I on the Ca2+ regulation of ATP-binding and ADP dissociation of myosin II in native cardiac myofibrils. Biochemistry 55, 1887–1897 (2016).

22. J. C. Barbato, Q.-Q. Huang, M. M. Hossain, M. Bond, J.-P. Jin, Proteolytic N-terminal truncation of cardiac troponin I enhances ventricular diastolic function. J. Biol. Chem. 280, 6602–6609 (2005).

23. D. G. Ward, M. P. Cornes, I. P. Trayer, Structural consequences of cardiac troponin I phosphorylation. J. Biol. Chem. 277, 41795–41801 (2002).

24. H.-Z. Feng, M. Chen, L. S. Weinstein, J.-P. Jin, Removal of the N-terminal extension of cardiac troponin I as a functional compensation for impaired myocardial β-adrenergic signaling. J. Biol. Chem. 283, 33384–33393 (2008).

25. H.-Z. Feng, X. Huang, J.-P. Jin, N-terminal truncated cardiac troponin I enhances Frank-Starling response by increasing myofilament sensitivity to resting tension. J. Gen. Physiol. 155, e202012821 (2023).

26. C. M. Warren, M. Halas, H.-Z. Feng, B. M. Wolska, J.-P. Jin, R. J. Solaro, NH2-Terminal Cleavage of Cardiac Troponin I Signals Adaptive Response to Cardiac Stressors. J. Cell. Signal. 2 (2021).

27. A. E. Messer, A. M. Jacques, S. B. Marston, Troponin phosphorylation and regulatory function in human heart muscle: dephosphorylation of Ser23/24 on troponin I could account for the contractile defect in end-stage heart failure. J. Mol. Cell. Cardiol. 42, 247–259 (2007).

28. Y. Li, P.-Y. J. Charles, C. Nan, J. R. Pinto, Y. Wang, J. Liang, G. Wu, J. Tian, H.-Z. Feng, J. D. Potter, J.-P. Jin, X. Huang, Correcting diastolic dysfunction by Ca2+ desensitizing troponin in a transgenic mouse model of restrictive cardiomyopathy. J. Mol. Cell. Cardiol. 49, 402–411 (2010).

29. J. F. Shaffer, T. E. Gillis, Evolution of the regulatory control of vertebrate striated muscle: the roles of troponin I and myosin binding protein-C. Physiol. Genomics 42, 406–419 (2010).

30. M. Vornanen, Maximum heart rate of soricine shrews: correlation with contractile properties and myosin composition. Am. J. Physiol. 262, R842–851 (1992).

31. K. D. Jürgens, R. Fons, T. Peters, S. Sender, Heart and respiratory rates and their significance for convective oxygen transport rates in the smallest mammal, the Etruscan shrew Suncus etruscus. J. Exp. Biol. 199, 2579–2584 (1996).

32. M. Vornanen, Basic functional properties of the cardiac muscle of the common shrew (Sorex araneus) and some other small mammals. J. Exp. Biol. 145, 339–351 (1989).

33. Y. H. Chang, B. I. Sheftel, B. Jensen, Anatomy of the heart with the highest heart rate. J. Anat. 241, 173–190 (2022).

34. P. Morrison, F. A. Ryser, A. R. Dawe, Studies on the Physiology of the Masked Shrew Sorex cinereus. Physiol. Zool. 32, 256–271 (1959).

35. A. Nagel, The electrocardiogram of European shrews. Comp. Biochem. Physiol. A Physiol. 83, 791–794 (1986).

36. K. He, T. G. Eastman, H. Czolacz, S. Li, A. Shinohara, S. Kawada, M. S. Springer, M. Berenbrink, K. L. Campbell, Myoglobin primary structure reveals multiple convergent transitions to semi-aquatic life in the world’s smallest mammalian divers. eLife 10, e66797 (2021).

37. K. He, A. Shinohara, K. M. Helgen, M. S. Springer, X.-L. Jiang, K. L. Campbell, Talpid mole phylogeny unites shrew moles and illuminates overlooked cryptic species diversity. Mol. Biol. Evol. 34, 78–87 (2017).

38. L. Saggin, L. Gorza, S. Ausoni, S. Schiaffino, Troponin I switching in the developing heart. J. Biol. Chem. 264, 16299–16302 (1989).

39. U. Oron, M. Mandelberg, Comparative morphometry of the mitochondria and activity of some enzymes in the myocardium of small mammals. J. Mol. Cell. Cardiol. 17, 627–632 (1985).

40. A. Armsby, T. Quilliam, H. Soehnle, Some observations on the ecology of the mole. J. Zool. - J ZOOL 149, 110–112 (1966).

41. T. Allison, H. Van Twyver, Sleep in the moles, Scalopus aquaticus and Condylura cristata. Exp. Neurol. 27, 564–578 (1970).

42. K. L. Campbell, P. W. Hochachka, Thermal biology and metabolism of the American shrew-mole, Neurotrichus gibbsii. J. Mammal. 81, 578–585 (2000).

43. K. L. Campbell, I. W. McIntyre, R. A. MacArthur, Fasting metabolism and thermoregulatory competence of the star-nosed mole, Condylura cristata (Talpidae: Condylurinae). Comp. Biochem. Physiol. A. Mol. Integr. Physiol. 123, 293–298 (1999).

44. N. Blom, T. Sicheritz-Pontén, R. Gupta, S. Gammeltoft, S. Brunak, Prediction of post-translational glycosylation and phosphorylation of proteins from the amino acid sequence. Proteomics 4, 1633–1649 (2004).

45. G. Yeo, C. B. Burge, Maximum entropy modeling of short sequence motifs with applications to RNA splicing signals. J. Comput. Biol. J. Comput. Mol. Cell Biol. 11, 377–394 (2004).

46. P. J. Shepard, E.-A. Choi, A. Busch, K. J. Hertel, Efficient internal exon recognition depends on near equal contributions from the 3′ and 5′ splice sites. Nucleic Acids Res. 39, 8928–8937 (2011).

47. J.-J. Sheng, J.-P. Jin, Gene regulation, alternative splicing, and posttranslational modification of troponin subunits in cardiac development and adaptation: a focused review. Front. Physiol. 5, 165 (2014).

48. M. Rasmussen, J.-P. Jin, Troponin Variants as Markers of Skeletal Muscle Health and Diseases. Front. Physiol. 12 (2021).

49. Tissue expression of TNNI3 - Summary - The Human Protein Atlas. https://www.proteinatlas.org/ENSG00000129991-TNNI3/tissue.

50. F. M. Real, S. A. Haas, P. Franchini, P. Xiong, O. Simakov, H. Kuhl, R. Schöpflin, D. Heller, M.-H. Moeinzadeh, V. Heinrich, T. Krannich, A. Bressin, M. F. Hartmann, S. A. Wudy, D. K. N. Dechmann, A. Hurtado, F. J. Barrionuevo, M. Schindler, I. Harabula, M. Osterwalder, M. Hiller, L. Wittler, A. Visel, B. Timmermann, A. Meyer, M. Vingron, R. Jiménez, S. Mundlos, D. G. Lupiáñez, The mole genome reveals regulatory rearrangements associated with adaptive intersexuality. Science 370, 208–214 (2020).

51. J. R. Johnston, P. B. Chase, J. R. Pinto, Troponin through the looking-glass: emerging roles beyond regulation of striated muscle contraction. Oncotarget 9, 1461–1482 (2017).

52. S. Casas-Tintó, A. Maraver, M. Serrano, A. Ferrús, Troponin-I enhances and is required for oncogenic overgrowth. Oncotarget 7, 52631–52642 (2016).

53. C. Chen, J.-B. Liu, Z.-P. Bian, J.-D. Xu, H.-F. Wu, C.-R. Gu, Y. Shi, J.-N. Zhang, X.-J. Chen, D. Yang, Cardiac troponin I is abnormally expressed in non-small cell lung cancer tissues and human cancer cells. Int. J. Clin. Exp. Pathol. 7, 1314–1324 (2014).

54. L. S. McRobb, V. S. Lee, M. Simonian, Z. Zhao, S. G. Thomas, M. Wiedmann, J. V. A. Raj, M. Grace, V. Moutrie, M. J. McKay, M. P. Molloy, M. A. Stoodley, Radiosurgery Alters the Endothelial Surface Proteome: Externalized Intracellular Molecules as Potential Vascular Targets in Irradiated Brain Arteriovenous Malformations. Radiat. Res. 187, 66–78 (2017).

55. P. B. Chase, M. P. Szczypinski, E. P. Soto, Nuclear tropomyosin and troponin in striated muscle: new roles in a new locale? J. Muscle Res. Cell Motil. 34, 275–284 (2013).

56. J. R. Speakman, D. W. Thomas, “Physiological ecology and energetics of bats” in Bat Ecology, T. Kunz, M. Fenton, Eds. (University of Chicago Press, Chicago, 2003), pp. 430–490.

57. D. K. N. Dechmann, S. Ehret, A. Gaub, B. Kranstauber, M. Wikelski, Low metabolism in a tropical bat from lowland Panama measured using heart rate telemetry: an unexpected life in the slow lane. J. Exp. Biol. 214, 3605–3612 (2011).

58. M. T. O’Mara, M. Wikelski, C. C. Voigt, A. Ter Maat, H. S. Pollock, G. Burness, L. M. Desantis, D. K. Dechmann, Cyclic bouts of extreme bradycardia counteract the high metabolism of frugivorous bats. eLife 6, e26686 (2017).

59. C. Zhu, Z. Chen, W. Guo, Pre-mRNA mis-splicing of sarcomeric genes in heart failure. Biochim. Biophys. Acta 1863, 2056–2063 (2017).

60. S. E. Currie, G. Körtner, F. Geiser, Pronounced differences in heart rate and metabolism distinguish daily torpor and short-term hibernation in two bat species. Sci. Rep. 12, 21721 (2022).

61. M. T. O’Mara, S. Rikker, M. Wikelski, A. Ter Maat, H. S. Pollock, D. K. N. Dechmann, Heart rate reveals torpor at high body temperatures in lowland tropical free-tailed bats. R. Soc. Open Sci. 4, 171359 (2017).

62. S. M. Harrison, D. M. Bers, Influence of temperature on the calcium sensitivity of the myofilaments of skinned ventricular muscle from the rabbit. J. Gen. Physiol. 93, 411–428 (1989).

63. T. Veltri, G. A. P. de Oliveira, E. A. Bienkiewicz, F. L. Palhano, M. de A. Marques, A. H. Moraes, J. L. Silva, M. M. Sorenson, J. R. Pinto, Amide hydrogens reveal a temperature-dependent structural transition that enhances site-II Ca2+-binding affinity in a C-domain mutant of cardiac troponin C. Sci. Rep. 7, 691 (2017).

64. B. Liu, L. C. Wang, D. D. Belke, Effects of temperature and pH on cardiac myofilament Ca2+ sensitivity in rat and ground squirrel. Am. J. Physiol. 264, R104–8 (1993).

65. W. K. Milsom, M. B. Zimmer, M. B. Harris, Regulation of cardiac rhythm in hibernating mammals. Comp. Biochem. Physiol. A. Mol. Integr. Physiol. 124, 383–391 (1999).

66. J. Taylor, “Evolution of energetic strategies in shrews” (1998), pp. 309–346.

67. S. Engelhardt, L. Hein, F. Wiesmann, M. J. Lohse, Progressive hypertrophy and heart failure in β1-adrenergic receptor transgenic mice. Proc. Natl. Acad. Sci. 96, 7059–7064 (1999).

68. K. Gao, A. Masuda, T. Matsuura, K. Ohno, Human branch point consensus sequence is yUnAy. Nucleic Acids Res. 36, 2257–2267 (2008).

69. J. Xie, L. Wang, R.-J. Lin, Variations of intronic branchpoint motif: identification and functional implications in splicing and disease. Commun. Biol. 6, 1–5 (2023).

70. E. Lara-Pezzi, J. Gómez-Salinero, A. Gatto, P. García-Pavía, The Alternative Heart: Impact of Alternative Splicing in Heart Disease. J Cardiovasc. Transl. Res. 6, 945–955 (2013).

71. A. Beqqali, Alternative splicing in cardiomyopathy. Biophys. Rev. 10, 1061–1071 (2018).

72. M. Gotthardt, V. Badillo-Lisakowski, V. N. Parikh, E. Ashley, M. Furtado, M. Carmo-Fonseca, S. Schudy, B. Meder, M. Grosch, L. Steinmetz, C. Crocini, L. Leinwand, Cardiac splicing as a diagnostic and therapeutic target. Nat. Rev. Cardiol. 20, 517–530 (2023).

73. Z. Mahmud, S. Zahran, P. B. Liu, B. Reiz, B. Y. H. Chan, A. Roczkowsky, C.-S. E. McCartney, P. L. Davies, L. Li, R. Schulz, P. M. Hwang, Structure and proteolytic susceptibility of the inhibitory C-terminal tail of cardiac troponin I. Biochim. Biophys. Acta Gen. Subj. 1863, 661–671 (2019).

74. T. G. Martin, J. A. Kirk, Under construction: The dynamic assembly, maintenance, and degradation of the cardiac sarcomere. J. Mol. Cell. Cardiol. 148, 89–102 (2020).

75. J. Kühnisch, C. Herbst, N. Al-Wakeel-Marquard, J. Dartsch, M. Holtgrewe, A. Baban, G. Mearini, J. Hardt, K. Kolokotronis, B. Gerull, L. Carrier, D. Beule, S. Schubert, D. Messroghli, F. Degener, F. Berger, S. Klaassen, Targeted panel sequencing in pediatric primary cardiomyopathy supports a critical role of TNNI3. Clin. Genet. 96, 549–559 (2019).

76. R. W. Meredith, J. E. Janečka, J. Gatesy, O. A. Ryder, C. A. Fisher, E. C. Teeling, A. Goodbla, E. Eizirik, T. L. L. Simão, T. Stadler, D. L. Rabosky, R. L. Honeycutt, J. J. Flynn, C. M. Ingram, C. Steiner, T. L. Williams, T. J. Robinson, A. Burk-Herrick, M. Westerman, N. A. Ayoub, M. S. Springer, W. J. Murphy, Impacts of the Cretaceous Terrestrial Revolution and KPg extinction on mammal diversification. Science 334, 521–524 (2011).

77. S. F. Altschul, T. L. Madden, A. A. Schäffer, J. Zhang, Z. Zhang, W. Miller, D. J. Lipman, Gapped BLAST and PSI-BLAST: a new generation of protein database search programs. Nucleic Acids Res. 25, 3389–3402 (1997).

78. M. J. Sullivan, N. K. Petty, S. A. Beatson, Easyfig: a genome comparison visualizer. Bioinforma. Oxf. Engl. 27, 1009–1010 (2011).

79. J. O. Wertheim, B. Murrell, M. D. Smith, S. L. Kosakovsky Pond, K. Scheffler, RELAX: Detecting relaxed selection in a phylogenetic framework. Mol. Biol. Evol. 32, 820–832 (2015).

80. S. L. K. Pond, S. D. W. Frost, S. V. Muse, HyPhy: hypothesis testing using phylogenies. Bioinformatics 21, 676–679 (2005).

81. R. W. Meredith, J. Gatesy, W. J. Murphy, O. A. Ryder, M. S. Springer, Molecular decay of the tooth gene enamelin (ENAM) mirrors the loss of enamel in the fossil record of placental mammals. PLOS Genet. 5, e1000634 (2009).

82. A. J. Sabucedo, K. G. Furton, Estimation of postmortem interval using the protein marker cardiac Troponin I. Forensic Sci. Int. 134, 11–16 (2003).

83. M. S. Utter, C. M. Warren, R. J. Solaro, Impact of anesthesia and storage on posttranslational modifications of cardiac myofilament proteins. Physiol. Rep. 3 (2015).

84. J. Kütt, G. Margus, L. Kask, T. Rätsepso, K. Soodla, R. Bernasconi, R. Birkedal, P. Järv, M. Laasmaa, M. Vendelin, Simple analysis of gel images with IOCBIO Gel. BMC Biol. 21, 225 (2023).

85. Y. Wang, Y. Zhang, W. Hu, S. Xie, C.-X. Gong, K. Iqbal, F. Liu, Rapid alteration of protein phosphorylation during postmortem: implication in the study of protein phosphorylation. Sci. Rep. 5, 15709 (2015).

86. C. Natarajan, X. Jiang, A. Fago, R. E. Weber, H. Moriyama, J. F. Storz, Expression and Purification of Recombinant Hemoglobin in Escherichia coli. PLOS ONE 6, e20176 (2011).

