## Supplementary Material for "Genetic excision of the regulatory cardiac troponin I extension in high heart rate mammal clades"

### Supplementary Materials

#### Materials and Methods

##### 1. Experimental subject details

Samples were acquired as described in table S3.

**Table S3.** Origins of samples used in this study. * Permission for collection of all samples collected in China, Vietnam and Japan was obtained from all the local forest bureaus or nature reserve administrations.

| Latin name | Common name | Source | Other notes | Experiment |
|---|---|---|---|---|
| *Erinaceus europaeus* | West European hedgehog | Private veterinary practice (United Kingdom). | Female juvenile euthanised by intrahepatic pentobarbital by veterinary surgeon. Animal had severe pneumonia secondary to lung worm infection and showed deteriorating health despite appropriate treatment with antiparaciticides (moxidectin/imidacloprid), antibiotic (enrofloxacin) and steroid (dexamethasone). Heart sample collected ~2 hr post mortem. Heart preserved in RNAlater and stored at -80ºC. | Western blot, mRNA sequencing |
| *Crocidura russula* | Greater white-toothed shrew | Killed by domestic cat (Valencia, Spain). | Unconfirmed time of death, likely >24 hr but not yet showing signs of decay. Estimated as 24 – 72 hr post mortem.<br>Heart preserved in RNAlater and stored at -80ºC. | Western blot, mRNA sequencing |
| *Sorex minutus* | Eurasian pygmy shrew | Killed by domestic cat (West Yorkshire, United Kingdom). | Two samples; one estimated to be <12 hr post mortem, one <1 hr post mortem.<br>Heart preserved in RNAlater and stored at -80ºC. | Western blot |
| *Mus musculus* | House mouse (laboratory mouse). | Holly Shiels laboratory (University of Manchester, United Kingdom).<br><br>Katinka Stecina laboratory (University of Manitoba, Winnipeg, Canada).<br><br>Guangdong Medical Laboratory animal center (Guangzhou, China). | Figure 1B, S2A,B: C57BL/6 strain, male, 10 weeks old, wild-type (no genetic modifications).<br>Sacrificed with cervical dislocation by trained personnel according to Home Office Schedule 1 regulations and with institutional approval. Heart snap-frozen and stored at -80ºC.<br><br>Figure 1C: C57BL/6, female, 16 week-old Sim1CreTdTom-ChR2 mouse, anaesthetized with isoflurane and heart surgically dissected out for protein extraction and storage at -65ºC.<br><br>Figure 4: C57BL/6 strain, female, 8 weeks old, wild-type (no genetic modifications) | Western blot |
| *Sorex palustris* | American water shrew | Nopiming Provincial Park, Manitoba, Canada. | Two animals were captured under permission of a Government of Manitoba Conservation trapping permit (WB24532), and cared for in accordance with the principles and guidelines of the Canadian Council of Animal Care (University of Manitoba Animal Use Protocol: F20-019 (AC11607)). Heart stored at -80ºC. | mRNA sequencing |
| *Sorex cinereus* | Masked shrew | Winnipeg, Manitoba, Canada. | Three animals were captured under permission of a Government of Manitoba Conservation trapping permit (WB24532), and cared for in accordance with the principles and guidelines of the Canadian Council of Animal Care (University of Manitoba Animal Use Protocol: F20-019 (AC11607)). Heart stored at -80ºC. | mRNA sequencing |
| *Galemys pyrenaicus* | Pyrenean desman | Jorge Gonzales Esteban (DESMA | Five samples from desmans that died from accidental/natural causes and were frozen at -20ºC for between 3 months and 12 years. | Western blot, mRNA sequencing, |

| | | | | |
|---|---|---|---|---|
| | | Estudios Ambientales, Spain) | Frozen animals were dissected and hearts then preserved in RNAlater and protein extracts stored at -80ºC. | mass spectrometry analysis |
| *Talpa europaea* | European mole | Professional molecatcher using mechanical traps (West Yorkshire, United Kingdom). | Two samples; both <18 hr post-mortem and exhibiting rigor mortis.<br>Heart preserved in RNAlater and stored at -80ºC. | Western blot, mRNA sequencing |
| *Desmana moschata* | Russian desman | Anna Bannikova | Tissue sample. Previously used (*36*, *37*) | DNA sequencing |
| *Uropsilus gracilis* | Gracile shrew-like mole | Kai He | Tissue sample. Previously used (*36*, *37*) | DNA sequencing |
| *Urotrichus talpoides* | Greater Japanese shrew mole | Shin-ichiro Kawada | Tissue sample. Previously used (*36*, *37*) | DNA sequencing |
| *Anourosorex squamipes* | Chinese mole shrew | Kai He | One specimen captured at Mt. Ailaoshan, Yunnan, China. Heart tissue preserved in RNAlater, and stored at -80ºC before sequencing.<br>One specimen captured at Chengdu, China. Heart tissue was frozen in liquid nitrogen and stored at -80ºC before western blotting.* | Transcriptome sequencing, western blot |
| *Blarina brevicauda* | Northern short-tailed shrew | Kevin Campbell | Previously collected specimen obtained following University of Manitoba Animal Use Protocol F05-014 and stored at -20°C. * | Mass-spectrometry analysis |
| *Blarinella quadraticauda* | Asiatic short-tailed shrew | Kai He | One specimen captured at Mt. Ailaoshan, Yunnan, China. Heart tissue preserved in RNAlater, and stored at -80ºC. * | Transcriptome sequencing |
| *Chodsigoa smithii* | | Kai He | One specimen captured at Mt. Jinfoshan, Chongqing, China. Heart tissue preserved in RNAlater, and stored at -80ºC.* | Transcriptome sequencing |
| *Crocidura tanakae* | Taiwanese gray shrew | Kai He | One specimen captured at Mt. Chebaling, Guangdong China. Heart tissue preserved in RNAlater, and stored at -80ºC. * | Transcriptome sequencing |
| *Dymecodon pilirostris* | True's shrew mole | Kai He | One specimen captured at Mt. Nishikoma, Nagano, Japan. Heart tissue preserved in RNAlater, and stored at -80ºC. * | Transcriptome sequencing |
| *Episoriculus umbrinus* | Hidden brown-toothed shrew | Kai He | One specimen captured at Mt. Ailaoshan, Yunnan, China. Heart tissue preserved in RNAlater, and stored at -80ºC. * | Transcriptome sequencing |
| *Nectogale elegans* | Elegant water shrew | Kai He | One specimen captured at Mt. Wuliangshan, Yunnan, China. Heart tissue preserved in RNAlater, and stored at -80ºC. * | Transcriptome sequencing |
| *Neotetracus sinensis* | Shrew gymnure | Kai He | One specimen captured at Mt. Ailaoshan, Yunnan, China. Heart tissue preserved in RNAlater, and stored at -80ºC. * | Transcriptome sequencing |
| *Sorex bedfordiae* | Lesser striped shrew | Kai He | One specimen captured at Mt. Ailaoshan, Yunnan, China. Heart tissue preserved in RNAlater, and stored at -80ºC. * | Transcriptome sequencing |
| *Sorex hosonoi* | Azumi shrew | Kai He | One specimen captured at Mt. Nishikoma, Nagano, Japan. Heart tissue preserved in RNAlater, and stored at -80ºC. * | Transcriptome sequencing |
| *Chimarrogale himalayica varennei* | Himalayan water shrew | Masaharu Motokawa | Two specimens captured at Vietnam. Heart tissue preserved in RNAlater, and stored at -80ºC. * | Transcriptome sequencing |
| *Uropsilus gracilis* | Gracile shrew-like mole | Kai He | One specimen captured at Mt. Jinfoshan, Chongqing, China. Heart tissue preserved in RNAlater, and stored at -80ºC. * | Transcriptome sequencing |
| *Urotrichus talpoides* | Greater Japanese shrew mole | Kai He | One specimen captured at Mt. Nishikoma, Nagano, Japan. Heart tissue preserved in RNAlater, and stored at -80ºC. * | Transcriptome sequencing |
| *Suncus murinus* | House shrew | Kai He | One specimen captured at Guangzhou, China. Heart tissue was frozen in liquid nitrogen and stored at -80ºC before western blot. * | Western blot, mass spectrometry analysis |
| *Hipposideros armiger* | Great roundleaf bat | Kai He | Two specimens captured at Huizhou, China. Heart tissue was frozen in liquid nitrogen and stored at -80ºC before western blot. * | Western blot, mass spectrometry analysis |

#### 2. *TNNI3*/cTnI nucleotide/protein sequences

##### *2.1 Gene predictions and genome annotation*

Gnomon-predicted *TNNI3* coding sequences from three eulipotyphlan species (*Sorex araneus*, *Condylura cristata*, and *Erinaceus europaeus*) were first retrieved from National Center for Biotechnology Information (NCBI) GenBank using key word searches and the *TNNI3* ortholog page (https://www.ncbi.nlm.nih.gov/gene/7137/ortholog/?scope=8782&term=TNNI3). As the *C. cristata* sequence (GB accession number XM_004694091.2) was found to be mis-annotated, the contig housing the gene (AJFV01074313.1) was downloaded and manually annotated. The resulting coding sequences were used as nucleotide blast (blastn) (*77*) queries against eulipotyphlan whole-genome shotgun sequences using the "discontinuous megablast" setting. Contigs returned as hits were downloaded and annotated using the Geneious (version 9.1.8) annotate function. In cases where gaps in coding sequence were apparent (e.g., *Uropsilus nivatus, Crocidura indochinensis*) or for sequenced species for which whole-genome shotgun sequences were unavailable (i.e., *Neohylomys hainanensis*), corresponding regions from closely related species were used as blastn queries against sequence read archive (SRA) files. All hits were downloaded and assembled to coding regions of closely related species using the Geneious map to reference function. As only raw (unassembled) genomic data were available for *S. fumeus* (SRR18739258), SRA files for this species were first converted to fastq sequence reads using the *fastq-dump* function in the SRA-Toolkit v3.0.0 (NCBI). Fastq reads were trimmed of adapter and low-quality sequences using BBDuk (Joint Genomics Institute) with the flags mink=11, hdist=1, tpe, tbo, qtrim=rl and timq=10. Trimmed reads were assembled to a *TNNI3* reference sequence (*Sorex palustris*) using BBMap (Joint Genomics Institute) with the flags maxindel=80 and k=10. All mined data were imported to Geneious and annotated as above. Linear sequence comparison figures of the genomic data were prepared using EasyFig (*78*).

##### *2.2 Transcriptomics and mRNA sequencing*

Heart samples collected from 13 eulipotyphlan species were placed in RNAlater (Qiagen, China) and stored at -80°C. Total RNA was extracted from each specimen and *de novo* assembled transcriptomes were generated using Trinity v2.4.1. with default parameters as previously described (*36*). The species list and source of tissues for mRNA extraction are provided under '1. Experimental subject details'.

Total mRNA was also extracted from ~60 mg of heart tissue of five eulipotyphlan species (table S3) using Trizol (Invitrogen) following the manufacturer's instructions. cDNA was generated with SMARTScribe Reverse Transcriptase (Takara) or ProtoScript II Reverse Transcriptase (NEB) according to the manufacture's recommendations. PCRs were performed using Phusion DNA Polymerase with GC buffer supplemented with 3% DMSO. Thermocycling conditions followed the manufacture's recommendations, with the exception of annealing temperature and cycle count, which was optimised for each primer pair (table S3). Primers were designed from the predicted *TNNI3* sequences for a given species or closest available phylogenetic relatives. Amplicons of the expected size (~600 bp) were extracted from a 1.2% agarose gel in 1X SB Buffer and purified using GENEJet Gel Extraction Kit (Thermo Scientific). PCRs yielding non-specific amplification products were used as template in a nested PCR, using the primers defined in table S3, and then purified as above. Purified PCR amplicons were sequenced in both directions using the amplification primers by either Eurofins Genomics or the

Toronto Sick Kids Centre for Applied Genomics. All new sequences have been deposited in GenBank (see accession numbers in data S1).

Publicly deposited heart transcriptome SRA data from 11 chiropteran and five additional eulipotyphlan species (see table S1) were also downloaded to Geneious and assembled to both *TNNI3* and *TNNI1* coding sequences that we annotated for these same or closely related species using the "Map to Reference" function. To test for alternative exon 3 splicing in the 12 species with intact exon 3 sequences, SRA reads were first assembled to manually modified mRNA transcripts lacking exon 3. The resulting assemblies failed to provide evidence for alternative exon 3 splicing in both erinaceid (*Atelerix albiventris*) heart transcriptomes, though provided unequivocal evidence for alternative exon 3 splicing in all 10 available *Artibeus*, *Hipposideros*, and *Rhinolophus* specimens. To estimate the relative expression levels of alternatively spliced (lacking exon 3) and full-length (possessing exon 3) *TNN13* transcripts for the latter 10 bat specimens, 66 bp contigs (containing only exons 1, 2 and 4, and only exons 1, 2 and 42 bp of exon 3, respectively) were first constructed and used as templates for the SRA assemblies (maximum 5% mismatch threshold for both). A similar strategy was used to estimate the relative expression levels of *TNNI1* and *TNNI3* for the above 16 species, whereby coding sequences of *TNNI1* exons 3-7 and *TNNI3* 4-8 were used as templates for the SRA assemblies (maximum 10% mismatch threshold for both). All assemblies were visually inspected to ensure negligible off-target sequences were incorporated into the assemblies.

##### *2.3 PCR of exon 3 region*

As sequence data spanning the exon 3 region of *TNNI3* were unavailable for one of the major mole clades (shrew moles), we used previously extracted *Urotrichus talpoides* DNA (*36*, *37*) to PCR amplify this region. Primers in the upstream UTR and exon 4 were designed from known mRNA (transcriptome) sequences of this species. A similar strategy was employed to target this region from previously extracted *Uropsilus gracilis* DNA (*36*, *37*). Primers designed from the upstream UTR and exon 5 of the Pyrenean desman were also used to amplify the N-terminal region from previously purified Russian desman (*Desmana moschata*) DNA (*36*, *37*). PCRs were performed, and the resulting amplicons (700-1100 bp) were purified and sequenced as described in 1.2.

**Table S4.** List of primers and polymerize chain reaction conditions to amplify *TNNI3* from select eulipotyphlan species.

| Sample | Template | Forward primer | Reverse primer | Annealing temperature (°C) | Number of cycles |
|---|---|---|---|---|---|
| *Sorex palustris* | mRNA (cDNA) | TTTCTCTGCCTCCACAGCTC | CCTCAAACTTCTTCTTGCGG | 55 | 30 |
| *Sorex cinereus* | mRNA (cDNA) | TTTCTCTGCCTCCACAGCTC | CCTCAAACTTCTTCTTGCGG | 55 | 30 |
| *Talpa europaea* | mRNA (cDNA) | ATGGCGGACGAGAGCAGAGA | CCCTCGAACTTCTTCTTGCGG | 66.1 | 30 |
| *Erinaceus europaeus* | mRNA (cDNA) | TGGAACTCCTGATCTGATCG | TTTATTCCACAGGACCCTCTGG | 60 | 30 |
| *Galemys pyrenaicus* | mRNA (cDNA) | TTCCTCAGGCGCCGAAACG | CCTGCATCTTCGGGTCTGAGG | 60.6 | 30 |

| | | | | | |
|---|---|---|---|---|---|
| *Desmana moschata* | DNA | ATATTTAGTCTCTGTCCTGCC<br>CACGGAGCCTCACGCCAAGGT | GCTGCAGCTTTCTTGAGG<br>TGGGCCACCTGCAGCATCAG | 58.4 | 30 |
| *Uropsilus gracilis* | DNA | ATATTTAGTCTCTGCCCTCGCC<br>AGGAGCAGAGATGCGGTGAG | TCAGCTGCAGTTTCCTTGAG<br>TCCTTGAGGCCGAGATCTTAGG | 70.0<br>70.0 | 30<br>30 |
| *Urotrichus talpoides* | DNA | AGGAGCAGAGAAGCGGTGAG | TTCTTGAGGCGGAGATCTTAGAC | 60.6 | 30 |

#### 2.4 Selection analyses

Selection analyses were performed with the codeml program in PAML 4.9 (*35*) and the RELAX program (*79*) of HyPhy (*80*). Analyses were performed with (a) complete protein-coding sequences of the *TNNI3* gene (except for exon 3) for 48 placental mammal taxa (codeml and RELAX) and (b) exon 3 sequences for a reduced data set with 32 taxa that did not include shrews (RELAX only as codeml failed to return results; see data S2). The 48 placental mammal taxa include representatives from all four placental superorders (i.e., Afrotheria, Xenarthra, Laurasiatheria, and Euarchontoglires), but have enriched sampling within Laurasiatheria (data S1). Analyses with codeml were performed with two different codon frequency models, CodonFreq = 1 and CodonFreq = 2. We used cleandata = 0 (sites with ambiguity data were not removed) and method = 0 (dN/dS [=ω] values were estimated simultaneously across the tree). First, we compared a model with one dN/dS category for all branches to a model with two dN/dS branch categories for the protein-coding sequence alignment that included exons 1, 2, and 4-8. (dN/dS is the ratio of nonsynonymous to synonymous substitution rates.) In the two-category branch model, the first category was for branches leading to taxa with an intact copy of the coding sequence for exon 3 and the second category was for branches with one or more inactivating mutations in exon 3 (crown *Talpa* + stem *Talpa* + *Condylura* + *Urotrichus* + crown *Uropsilus* + stem *Uropsilus* + crown Soricidae + stem Soricidae). We performed these analyses to determine if exons 1, 2, and 4-8 have remained under purifying selection in taxa with a pseudoexonic copy of exon 3. We used the chi2 program in PAML to perform a log-likelihood ratio test on models with one versus two branch categories. We also performed an analysis with nine different branch categories (background + crown *Talpa* + stem *Talpa* + *Condylura* + *Urotrichus* + crown *Uropsilus* + stem *Uropsilus* + crown Soricidae + stem Soricidae) to ascertain the range of dN/dS values on branches of the tree that are transitional or fully pseudoexonic (*sensu* (*81*)) for exon 3. Analyses with RELAX were performed with two different models. The first model (RELAX null) allowed for a single branch category. The second model (RELAX alternative) allowed for two branch categories: foreground (Test), background (Reference). Both models allowed for three different ω values within each branch category ($\omega_1$, $\omega_2$, $\omega_3$). Comparisons between the alternative and null models were performed with lnL ratio tests in RELAX (*79*). For the analysis with 48 complete coding sequences (minus exon 3), the foreground branch category included all branches that are transitional or fully pseudoexonic for exon 3, and the background branch category included all of the remaining branches. For exon 3, the data set included 32 taxa and we performed two analyses. In the first analysis, the foreground category included branches for six non-desman talpids (*Condylura cristata, Scalopus aquaticus, Talpa occidentalis, Uropsilus gracilis, Uropsilus nivatus, Urotrichus talpoides*) and the stem branch for *Uropsilus*, all of which have inactivating mutations for exon 3. All of the non-talpid branches were assigned to the background branch category and all of the remaining talpid branches were left as unclassified and therefore ignored in the ω calculations and statistical tests.

In the second analysis, both desmans (*Galemys pyrenaicus* and *Desmana moschata*) and the stem desman branch were assigned to the foreground branch category (both desmans have intact copies of exon 3). All of the remaining talpid branches were left as unclassified and all non-talpid branches were assigned to the background branch category. We performed this test to determine if the exon 3 sequences in the desmans have evolved under relaxed selection relative to other mammals even though the exon sequences remain intact. For both of the analyses with 32 taxa, premature stop codons were replaced with Ns, frameshift insertions were deleted, and frameshift deletions were filled in with Ns (frameshift deletions) in talpid taxa with inactivating mutations.

##### *2.5 Phosphorylation site scores*

Phosphorylation site scoring was performed using the NetPhos 3.1 (*44*) online server (https://services.healthtech.dtu.dk/services/NetPhos-3.1/). cTnI protein sequence for *G. pyrenaicus* (including the predicted peptide encoded by exon 3) were uploaded along with *E. europaeus, M. musculus and H. sapiens* sequences. PKA phosphorylation site scores were recorded for $Ser_{23/24}$ for all species. Scores above 0.500 are ‘positive’ indicating a residue is a target for a given kinase.

##### *2.6 Splice site likelihood modelling*

To investigate the potential for alternative splicing in desmans (*G. pyrenaicus* and *D. moschata*; these species have intact splice sites and, unlike other moles and shrews, no inactivating indels), we performed maximum entropy analysis of 3’ and 5’ splice site strength (*45*). For each exon, the 3’ splice site strength analysis was performed at the 3’ end of the preceding (upstream) intron and the 5’ splice site analysis was performed with the 5’ end of the subsequent (downstream) intron (*46*). We compared the desmans with several representative species in our study that also have intact exon 3 splice sites and no indels (*Homo sapiens*, human; *Mus musculus*, house mouse; *Solenodon paradoxus*, Hispaniolan solenodon; *Erinaceus europaeus*, European hedgehog; *Neohylomys hainanensis*, Hainan gymnure). Two bat species shown to exhibit alternative exon 3 splicing (*Hipposideros armiger,* great roundleaf bat; *Artibeus jamaicensis*, Jamaican fruit bat) were also included in this analysis. Calculations were performed using the server hosted on the Burge Lab website (http://hollywood.mit.edu/burgelab/maxent/Xmaxentscan_scoreseq_acc.html). Because this analysis is optimized for comparison with the human genome, we also made equivalent comparisons for the neighboring exons (exons 2 and 4) in *TNNI3* for each species to ensure that the output yielded consistent comparisons across species.

#### 3. Protein expression characterisation

##### 3.1 Western blotting

The species list and source of tissues are provided in table S3. Previous work has shown that although cardiac troponin I gradually degrades after death, producing lower molecular weight fragments, an intact cTnI band remains visible in western blots from mammalian heart tissues collected up to at least four days post mortem (*82*). Thus, hearts from several free-ranging species that were opportunistically collected within this timeframe were included in these experiments. Note that post-mortem times of <24 hr were verified in all species except the greater white-toothed shrew, *Crocidura russula*, for which time of death was unknown but estimated to be 24 to 72 h. To confirm that both exon $3^+$ and exon $3^-$ cTnI isoforms were incorporated into cardiac myofilaments, myofibrillar proteins were also isolated for the house mouse (*Mus musculus*), great roundleaf bat (*Hipposideros armiger*; 2 specimens), Asian house shrew (*Suncus murinus*), and Chinese mole shrew (*Anourosorex squamipes*) using a previously published protocol (*83*).

Protein extraction was performed by homogenization of cardiac ventricular tissues in RIPA buffer (ThermoFisher 89900 or Beyotime P0013B) with 1% protease and phosphatase inhibitor (PPI) cocktail (Sigma-Aldrich PPC1010 or Beyotime P1045). Protein concentration was measured with bicinchoninic acid (BCA) assay (Millipore 71285-3 or Sangon C503021-0500) and samples were diluted to 2 µg protein $\mu l^{-1}$ with RIPA buffer and PPIs, and were then denatured with an equal volume of 2x Laemmli buffer (Sigma-Aldrich S3401) and heating to 95°C for 5 min.

Proteins were separated using sodium dodecyl sulfate–polyacrylamide gel electrophoresis (SDS-PAGE). Samples were loaded (15 µg protein per lane) alongside protein ladders (5 µl per lane, BLUeye Prestained Protein Ladder, Sigma-Aldrich 94964) on tris-glycine gels (16% acrylamide 1.0 mm, ThermoFisher XP00165BOX or XP00160BOX) using an Invitrogen XCell SureLock Mini-Cell or a SINSAGE-PR2. Proteins were thereafter transferred to a PVDF membrane (Invitrolon PVDF/Filter Paper Sandwiches, 0.45 µm, ThermoFisher LC2005 or Beyotime FFP36) with an XCell II Blot Module or a SINSAGE-BLOT2. Total protein loading was visualised with No-Stain™ Protein Labeling Reagent (ThermoFisher A44717) following the manufacturer's instructions and imaged on a Bio-Rad ChemiDoc. Subsequently, blocking was performed for 1 hr at room temperature with 5% skimmed milk in tris-buffered saline with 0.05% Tween 20 (TBS-T).

cTnI western blotting was achieved with a mouse monoclonal primary antibody against troponin I (Santa-Cruz SC-133117, 1:1000 dilutions of the 200 µg $ml^{-1}$ stock) diluted in 2% skimmed milk in TBS-T which was incubated with the blot overnight at 4°C or for 1 hr at room temperature. After washing with TBS-T, a HRP-conjugated anti-mouse secondary antibody (Santa-Cruz sc-516102, 1:5000 dilution of 400 µg $ml^{-1}$ stock) in 2% skimmed milk TBS-T was used for a 1 hr incubation at room temperature. Blots were imaged on a Bio-Rad ChemiDoc and chemiluminescent signals were developed with Millipore/Immobilon Classico Western HRP substrate (Merck WBLUC0500). Densitometric analysis was performed using IOCBIO Gel (*84*).

We also compared protein kinase A (PKA) phosphorylated proteins after exogenous PKA treatment. The exogenous treatment was designed to account for rapid post mortem decay of native phosphorylation (*85*), which would otherwise have precluded comparisons between samples. One protein extract (40 µg per 50 µl reaction) from each species (mouse, West European hedgehog, greater white-toothed shrew, European pygmy shrew, Pyrenean desman, and European mole) was incubated with catalytic subunit of PKA (5 µl per reaction, New England Biolabs P6000SVIAL) in NEBuffer™ for Protein Kinases (New England Biolabs B6022SVIAL) with 200 µM adenosine triphosphate (New England Biolabs P0756S). The treatment was performed for 1 hr at 30°C and was terminated by addition of an equal volume of 2x Laemmli buffer (Sigma-Aldrich S3401) and heating to 95°C for 5 min. SDS-PAGE and western blotting was performed as described above, except 8 µg protein per lane was loaded. Antibodies designed to specifically detect N-terminal phosphorylated cTnI ($Ser_{23/24}$, i.e., in the region encoded by exon 3; Cell Signaling 4004S, rabbit primary antibody, 1:1000 dilution of unspecified stock concentration) or general phosphorylated protein kinase A (phospho-PKA) substrates (RRXS*/T*; New England Biolabs 9624S rabbit primary antibody, 1:1000 dilution of unspecified stock concentration). Incubation was performed for 1 hr at room temperature or 4°C overnight on shaker, respectively. This was followed by incubation with a HRP-conjugated goat anti-rabbit secondary antibody (New England Biolabs 7074P2; 1:3000 dilution of unspecified stock concentration) for 1 hr at room temperature. The blot was imaged as described above, then stripped using Restore Western Blot Stripping Buffer (ThermoFisher 21059) and the TnI antibody protocol was followed as described above.

##### 3.2 Ectopic expression of exon $3^+$ and exon $3^-$ desman cTnI proteins for western blot analysis

*Expression cassette construction.* Coding sequences encoding *Galemys pyrenaicus* cTnI proteins, both with and without exon 3, were optimized for expression in *E. coli* and synthesized *in vitro* by GeneArt (Regensburg, Germany). Gene cassettes were ligated into a custom expression vector (*86*) using a New England BioLabs Quick Ligation Kit as recommended by the manufacturer. Chemically competent JM109 (DE3) *E. coli* (Promega) were prepared using a Z-Competent *E. coli* Transformation Kit and Buffer Set (Zymo Research). cTnI expression vectors were transformed into JM109 (DE3) chemically competent *E. coli* according to manufacturer's recommendations, plated on LB agar containing ampicillin (100 µg/ml), and incubated for 18 hr at 37°C. A single colony from each transformation was cultured in 50 ml of LB broth for 16 hr at 37°C on an orbital shaker at 200 rpm. Post incubation, 5 ml of the culture was pelleted by centrifugation and plasmid DNA was isolated using a GeneJET Plasmid Miniprep Kit (Thermo Scientific). The plasmid sequence was verified by Sanger sequencing at The Centre for Applied Genomics (Toronto, Ontario). The remainder of the culture was supplemented with glycerol to a final concentration of 10% and stored at –80°C until needed for expression.

*Protein expression*. 45 ml of starter culture (above) was added to 2500 ml of TB media containing ampicillin (100 µg/ul) and Antifoam SE-15 (0.1%; Sigma) in a 3 L spinner flask. Cultures were grown in a circulating water bath at 37°C, stirred with a cross blade impeller at 800 rpm and supplemented with air (0.08 l/min). Growth was monitored until the culture absorbance at 600 nm reached 0.6–0.8. Expression was then induced by supplementing the media with 0.2 mM isopropyl β-D-1-thiogalactopyranoside and 20 g/L of glucose. The culture was incubated at 28°C for 16 hr and then cells were pelleted by centrifugation at 5000 x g for 10 min.

*Cell lysis*. Cell pellets were re-suspended in cell lysis buffer (33 mM Tris-HCl, 0.5 mM DTT; 3 ml per gram cell paste), supplemented with lysozyme (1 mg per gram cell paste) and mixed in an orbital shaker at 300 rpm for 20 min at 20°C. Samples were sonicated on ice using a Fisher Scientific model 705 sonic dismembrator at 32 W with 10 seconds on/20 seconds off cycles for 15 min. TnI protein expression was visualized with western blotting as described above.

##### 3.3. Immunoprecipitation and LC-MS/MS analysis of the 22 kDa *TNNI3* protein from desman and shrew heart lysates

To verify that the 22 kDa band corresponded to cTnI (as opposed to the product of another *TNNI* gene paralog), we immunoprecipitated the TnI proteins of Pyrenean desman (*Galemys pyrenaicus*) and northern short-tailed shrew (*Blarina brevicauda*) hearts using the anti-Troponin I (C-4) (sc-133117, Santa Cruz). Briefly, a 30 µl slurry of protein G-Sepharose beads (Pierce) was washed three times with 500 µl of RIPA buffer (50 mM Tris-HCl, pH 7.6, 150 mM NaCl, 1% Triton X-100, 0.5% deoxycholic acid, 0.1% SDS), and resuspended in 30 µl of PBS buffer

(MP Biomedicals). The beads were then incubated with 1.3 µl of the anti-Troponin I antibody and 100 µl of protein lysate at 4°C for 3 hr with rotation, and then washed three times each with 500 µl of RIPA buffer followed by elution with 15 µl of glycine buffer (0.15 M NaCl, 0.2 M glycine-HCl, pH 2.0) on ice for 5 min. The eluted proteins were denatured in SDS protein loading buffer at 95°C for 5 min and resolved in 4-20% SDS-polyacrylamide gradient gels. The gel was stained with Coomassie brilliant blue G250 for 1 hr, de-stained with 10% acetic acid, and the ~22 kDa band was visualized above a fluorescent light box and excised using a clean glass coverslip. Before mass spectrometry, the identity of the ~22 kDa bands were confirmed by western blot analysis using the anti-Troponin I antibody in a separate test. The *TNNI3* 22kDa bands were sent for LC-MS/MS mass spectrometry analysis at the Southern Alberta Mass Spectrometry (SAMS) Centre, University of Calgary. As protein coding sequences for both *TNNI1* (encoding ssTnI) and *TNNI3* (encoding cTnI) of these two species are either unavailable or misannotated on GenBank, they were provided to the SAMS facility and included in the resulting in-house analysis.

##### 3.4. Immunoprecipitation and targeted LC-MS/MS analysis of the 22 kDa *TNNI3* protein from bat and shrew heart lysates

Given that standard LC-MS/MS for protein identification (above) was unable to recover the *TNNI3* exon 2/exon 4 splicing boundary, we performed LC-MS/MS with parallel reaction monitoring to target the N-terminal region of the great roundleaf bat (*Hipposideros armiger*) and Asian house shrew (*Suncus murinus*). Heart lysates were immunoprecipitated similar to that described above, with the exception that we used 13 µL of primary antibody and 150 µL of protein lysate. The excised ~22 kDa bands were sent to Wininovate Bio (Guangzhou, China) for sequencing and the MS/MS data were analyzed with PEAKS Studio 8.5 (Bioinformatics Solutions Inc., Waterloo, Canada) with a local false discovery rate at peptide-spectrum matches as 1.0% after searching against target sequences database with a maximum of three missed cleavages. Acetylation (Protein N-term) were selected for variable modifications. Precursor and fragment mass tolerance were set to 10 ppm and 0.05 Da, respectively.

**Table S1.** Relative expression levels of *TNNI3* (encoding cTnI) and *TNNI1* (encoding ssTnI) for five eulipotyphlan and 7 chiropteran species. Expression levels for each gene were estimated by assembling publicly available heart transcriptomes for each species to exons 4-8 of *TNNI3* and exons 3-7 of *TNNI1*, respectively.

| Species | Accession # | Sex/age | *TNNI3* hits | *TNNI1* hits |
|---|---|---|---|---|
| **Eulipotyphla lacking *TNNI3* exon 3** | | | | |
| *Sorex fumeus* | SRR24416557 | Male; ~2 months | 66,004 (99.93%) | 43 (0.07%) |
| *Blarina brevicauda* | SRR17216480 | Unknown | 128,824 (99.99%) | 14 (0.01%) |
| *Blarina brevicauda* | SRR17216485 | Unknown | 91,832 (99.97%) | 32 (0.03%) |
| *Blarina brevicauda* | SRR17216490 | Unknown | 110,629 (99.99%) | 13 (0.01%) |
| *Blarina brevicauda* | SRR17216495 | Unknown | 70,094 (99.79%) | 148 (0.21%) |
| *Condylura cristata* | SRR17216500 | Unknown | 71,859 (99.99%) | 5 (0.01%) |
| *Scalopus aquaticus* | SRR17216400 | Unknown | 29,511 (99.24%) | 227 (0.76%) |
| *Scalopus aquaticus* | SRR17216405 | Unknown | 10,541 (96.65%) | 365 (3.35%) |
| **Eulipotyphla possessing *TNNI3* exon 3** | | | | |
| *Atelerix albiventris* | SRR12442734 | Male; adult | 10,630 (71.7%) | 4196 (28.3%) |
| *Atelerix albiventris* | SRR12442741 | Female; adult | 30,376 (96.91%) | 970 (3.09%) |
| **Chiroptera exhibiting alternative exon 3 splicing of *TNNI3*** | | | | |
| *Hipposideros armiger* | SRR1657902 | Female; adult | 48,370 (99.63%) | 182 (0.27%) |
| *Rhinolophus sinicus* | SRR2273739 | Male; adult | 70,249 (99.88%) | 87 (0.12%) |
| *Rhinolophus clivosus* | SRR13417526 | Male; adult | 67,669 (>99.99%) | 1 (<0.01%) |
| *Artibeus bogotensis* | SRR13417527 | Male; adult | 123,547 (>99.99%) | 6 (<0.01%) |
| *Artibeus jamaicensis* | SRR13417529 | Male; adult | 151,942 (99.95%) | 78 (0.05%) |
| *Artibeus jamaicensis* | SRR13417530 | Female; adult | 150,160 (99.78%) | 334 (0.22%) |
| *Artibeus jamaicensis* | SRR13417531 | Male; adult | 94,919 (99.51%) | 465 (0.49%) |
| *Artibeus jamaicensis* | SRR13417532 | Female; adult | 185,058 (99.86%) | 259 (0.14%) |
| *Artibeus planirostris* | SRR13417533 | Male; adult | 124,446 (99.96%) | 56 (0.04%) |
| *Artibeus fraterculus* | SRR13417534 | Male; adult | 101,826 (99.73%) | 278 (0.27%) |

**Table S2.** Relative expression levels of full-length (possessing exon 3; '*TNNI3* FL') and alternatively spliced (lacking exon 3; '*TNNI3* AS') *TNNI3* transcripts for seven chiropteran species exhibiting alternative exon 3 splicing. Expression levels for each transcript were estimated by assembling publicly available heart transcriptomes for each species to 66 bp contigs (containing only exons 1, 2 and 4, and only exons 1, 2 and 42 bp of exon 3, respectively).

| Species | Accession # | Sex/age | *TNNI3* FL hits | *TNNI1* AS hits |
|---|---|---|---|---|
| *Hipposideros armiger* | SRR1657902 | Female; adult | 7,611 (59.46%) | 5,190 (40.54%) |
| *Rhinolophus sinicus* | SRR2273739 | Male; adult | 9,711 (57.69%) | 7,121 (42.31%) |
| *Rhinolophus clivosus* | SRR13417526 | Male; adult | 9,155 (67.51%) | 4,406 (32.49%) |
| *Artibeus bogotensis* | SRR13417527 | Male; adult | 11,813 (49.08%) | 12,255 (50.92%) |
| *Artibeus jamaicensis* | SRR13417529 | Male; adult | 21,408 (65.58%) | 11,234 (34.42%) |
| *Artibeus jamaicensis* | SRR13417530 | Female; adult | 26,001 (64.10%) | 14,560 (35.90%) |
| *Artibeus jamaicensis* | SRR13417531 | Male; adult | 12,676 (62.09%) | 7,740 (37.91%) |
| *Artibeus jamaicensis* | SRR13417532 | Female; adult | 25,232 (69.09%) | 11,287 (30.91%) |
| *Artibeus planirostris* | SRR13417533 | Male; adult | 16,043 (65.00%) | 8,637 (35.00%) |
| *Artibeus fraterculus* | SRR13417534 | Male; adult | 12,720 (67.57%) | 6,105 (32.43%) |

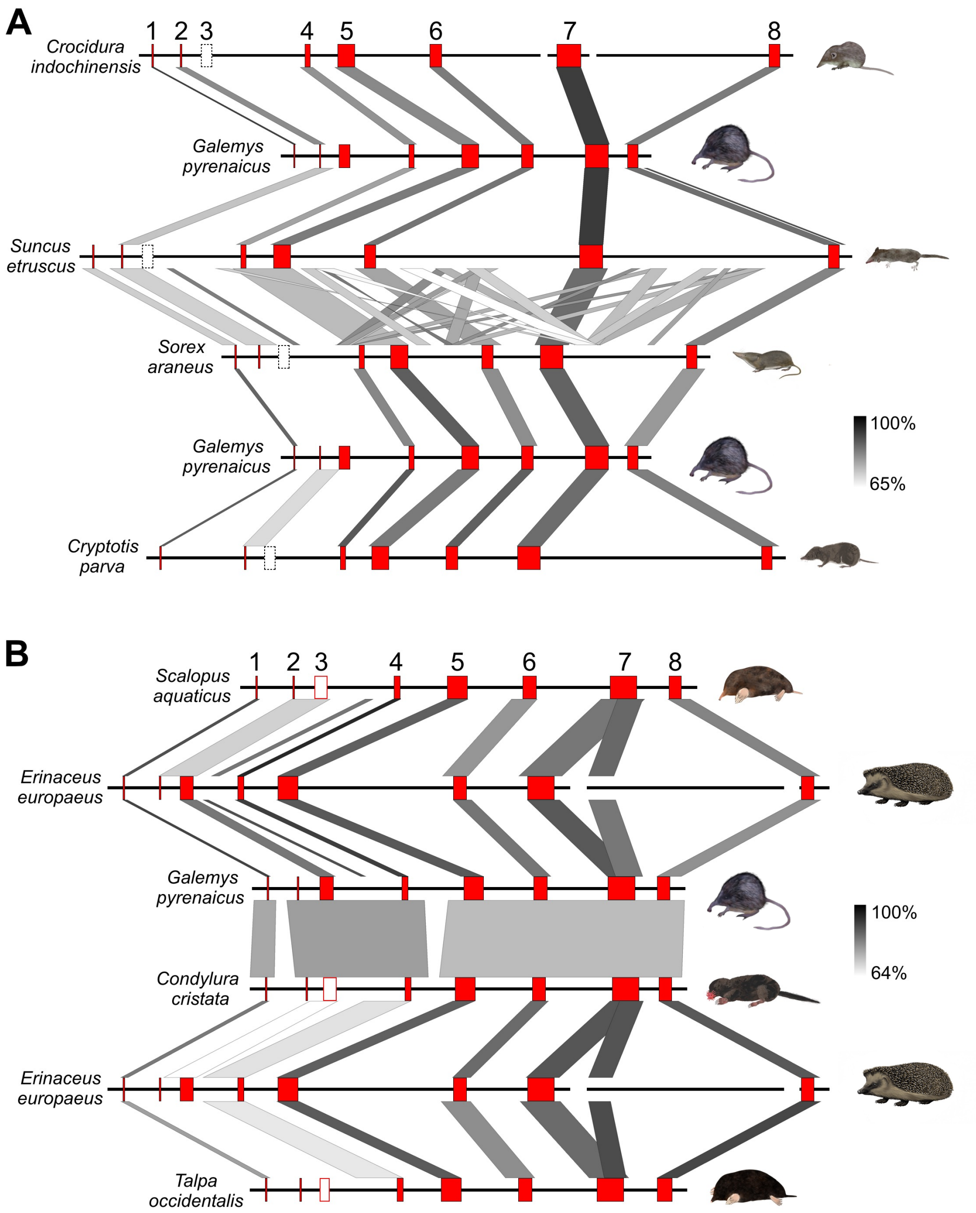

**Figure S1. Sequence identity comparisons of the *TNNI3* gene of select eulipotyphlan mammals.** (A) Linear comparisons illustrating the deletion of exon 3 (expected location denoted by open dashed boxes) in the genomes of four shrew species relative to the Pyrenean desman (*Galemys pyrenaicus*). B) Linear comparisons of four talpid mole species relative to the European hedgehog (*Erinaceus europaeus*). Intact exons are numbered and highlighted in red, while pseudoexonized (non-coding) exon 3 of moles is denoted by red open boxes. Thick black horizontal lines represent intron sequences while gaps denote missing data. Note that the sequence identity of the intact coding exons of moles (1, 2, 4-8) relative to corresponding coding exons of the hedgehog are higher than that of the non-coding exon 3 of moles, except for the Pyrenean desman whose intact exon 3 retains a relatively high sequence conservation.

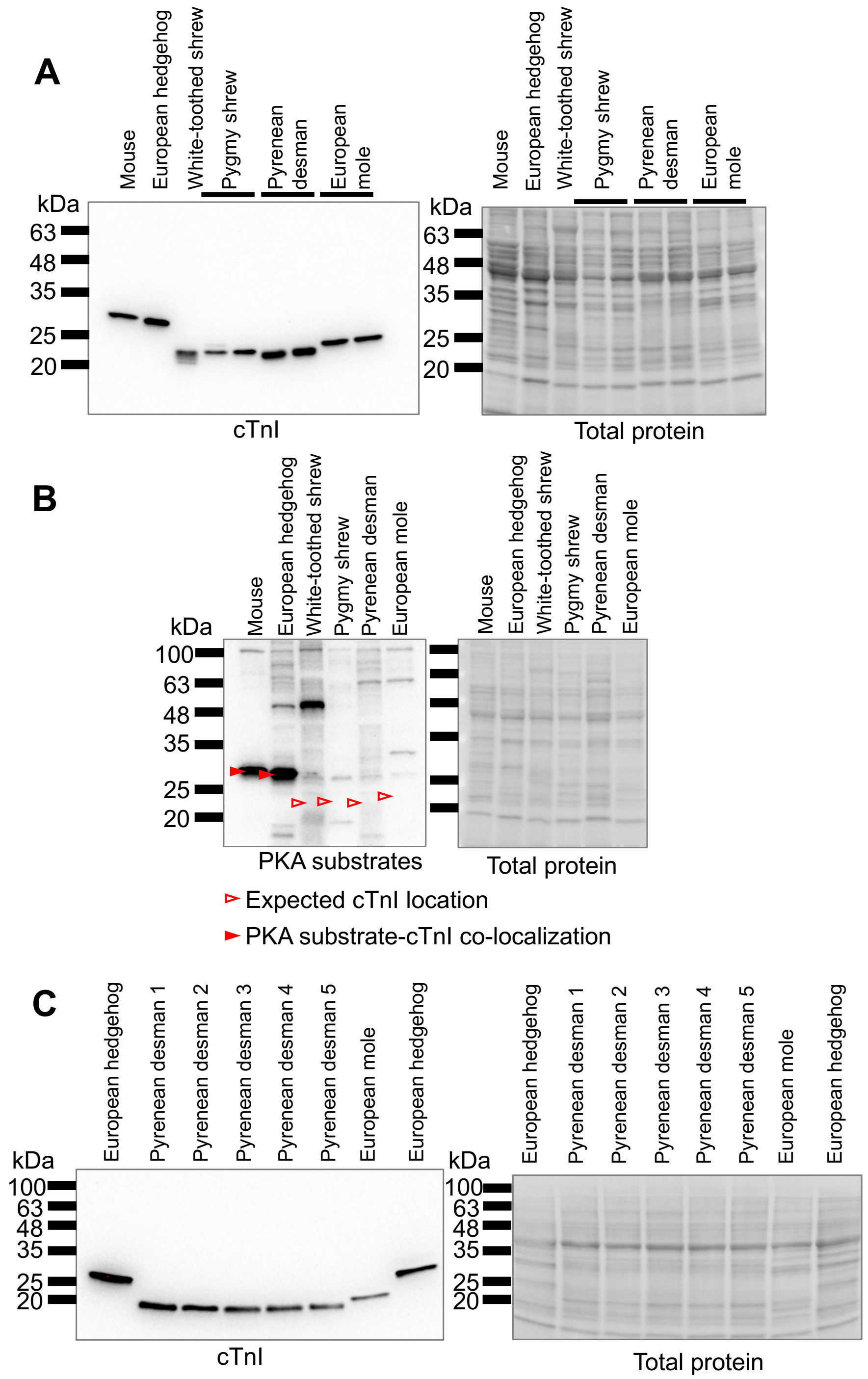

**Figure S2. Cardiac troponin I (cTnI) expression in heart extracts from representative eulipotyphlan species and mouse.** (A) cTnI western blotting including two individuals of pygmy shrew, two of five Pyrenean desmans (see panel C for all five), and two European moles. (B) Total PKA phosphorylation of protein kinase A treated cardiac extracts from five eulipotyphlans and mouse. PKA targets were detected with a primary antibody designed to identify phosphorylated serine (or threonine) with arginine at the -3 and -2 positions. This includes the cardiac troponin I (cTnI) N-terminal extension as well as other unidentified PKA substrates. Red arrows indicate the expected position of cTnI (filled arrows show co-localised PKA substrate). For a blot on aliquots of the same samples with a phosho-cTnI specific antibody, see Fig. 1B. (C) cTnI western blotting in all five desman heart samples available for the study. Note that the same hedgehog sample was used in the two flanking lanes to provide an N-terminal extension-present standard. Two Pyrenean desman extracts (4 and 5) and the European mole were the same as used in panel A. Total protein was visualised with No-Stain Protein Labeling Reagent.

***Blarina brevicauda***

Peptide matches to *Blarina* cTnI: **6 matches (4 non-duplicate, 2 duplicate)**

| Hit | Start-end | Observed mass | Expected mass | Calculated mass | Ion score | Expect value | Rank order | Peptide |
|---|---|---|---|---|---|---|---|---|
| 1 | 23-30 | 467.2777 | 932.5408 | 932.5365 | 26 | 0.0025 | 1 | K.TLMLQIAK.Q + Oxidation (M) |
| 2 | 93-103 | 609.3216 | 1216.6286 | 1216.6299 | 35 | 0.00031 | 1 | K.NVSEIADLTQK.V |
| 3 | 121-134 | 755.3737 | 1508.7328 | 1508.7327 | 15 | 0.321 | 1 | R.ISADAMMQALLGTR.A + 2 Oxidation (M) |
| 4 | 121-134 | 755.3737 | 1508.7328 | 1508.7327 | 77 | 1.8e-08 | 1 | R.ISADAMMQALLGTR.A + 2 Oxidation (M) |
| 5 | 121-134 | 755.3737 | 1508.7328 | 1508.7327 | 58 | 1.4e-06 | 1 | R.ISADAMMQALLGTR.A + 2 Oxidation (M) |
| 6 | 168-178 | 589.7769 | 1177.5392 | 1177.5397 | 15 | 0.034 | 1 | K.NIDALSGMEGR.K + Oxidation (M) |

>*Blarina brevicauda TNNI3* (cTnI)
MADERGAAKKSKITASRKLQLK**TLMLQIAK**QELEREVEERRGEKGRALSARCQPLELAGLGFAELQDLCRQLHAQ
VDKVDEERYDVEAKVTK**NVSEIADLTQK**VIDLRGKFKRPTLRRVR**ISADAMMQALLGTR**AKESLDLRANLKQVKK
EDTEKVKENREVGDWRK**NIDALSGMEGR**KKKFEG

Peptide matches to *Blarina* ssTnI: **none**

>*Blarina brevicauda TNNI1* (ssTnI)
MPEVERKSKITASRKLLLKSLMLAKAKECWEQEHEEREAEKARYLAERIPTLQTRGLSLSALQDLCRDLHAQVEV
VDEERYDIEAKCLHNTREIKDLKLKVLDLRGKFKRPPLRRVRVSADAMLRALLGSKHKVSMDLRANLKSVKKEDT
EKERPVEVGDWRKNVEAMSGMEGRKKMFDAAKTPVTQ

---

***Galemys pyrenaicus***

Peptide matches to *Galemys* cTnI: **2 matches (2 non-duplicate, 0 duplicate)**

| Hit | Start-end | Observed mass | Expected mass | Calculated mass | Ion score | Expect value | Rank order | Peptide |
|---|---|---|---|---|---|---|---|---|
| 1 | 93-103 | 609.3210 | 1216.6274 | 1216.6299 | 16 | 0.022 | 1 | K.NVSEIADLTQK.V |
| 2 | 166-176 | 589.7759 | 1177.5372 | 1177.5397 | 13 | 0.047 | 1 | K.NIDALSGMEGR.K + Oxidation (M) |

>*Galemys pyrenaicus TNNI3* (cTnI)
MADESREAKKSKISASRKLQLKTLMLQVAKQELEREAEERRGEKGRALSARCEPLELARLGFAELQDLCRQLHAR
VDKVDEERYDVEAKVTK**NVSEIADLTQK**ILDLRGKFKRPALRRVRISADAMMQALLGTRAKESMDLRANLKQVKK
EDTEKENREVGDWRK**NIDALSGMEGR**KKKFEG

Peptide matches to *Galemys* ssTnI: **none**

>*Galemys pyrenaicus TNNI1* (ssTnI)
MPEVERKSKITASRKLMLKSLMLAKAKECWEQEREEREAEKARYLAEHIPTLQTRGLSLRALQELCRDLHAKVEL
VDEERYDIEAKCLHNTREIKDLKLKVLDLRGKFKRPPLRRVRVSADAMLRALLGSKHKVSMDLRANLKSVKKEDT
EKERPVEVGDWRKNVEAMSGMEGRKKMFDAAKSPTSQ

---

**Example MS/MS spectra corresponding to peptide: NIDALSGMEGR**

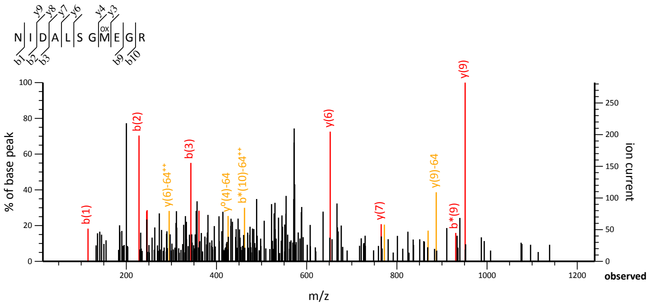

**Figure S3. Liquid chromatography with tandem mass spectrometry (LC-MS/MS) protein sequencing on the ~22 kDa cardiac TnI bands from a Pyrenean desman (*Galemys pyrenaicus*) and a northern short-tailed shrew (*Blarina brevicauda*).** Peptide matches to cTnI (*TNNI3*) are shown in red; no matches to ssTnI (*TNNI1*) were returned while peptide matches to other (non-TnI) protein products are not displayed.

```
Homo sapiens              ggtaagggctgggtggggtcttggtggtgatggggtccccactcctcctaaccc-agGCTAGGGAACCTC
Mus musculus              agggctggatg-ggagggtttgggtagtgatgtag-----cctgcttcaattcc-agGCTGGGGAACCGC
Solenodon paradoxus       ggggctgggta----------------acac---gtctctttaaccccaacccc-agGCGGGGGACCCAC
Erinaceus europaeus       ----ggacgaacgcgcgccgcgggatgacac----ggccgccgcccccgccccccagGCCGGGGAGCCCA
Neohylomys hainanensis    a--gaggactgggcagggtcttgggtgacac----taccgcgttccccgacccc-agGCCGGGGAGCCCT
Uropsilus gracilis*       ggcgctgatg-------------------------------------cagcccc-aaGCGGGGC-CCCGT
Uropsilus nivatus*        ggcgctgatg-------------------------------------cagcccc-aaGCGGGGC-CCCGT
Scalopus aquaticus*       ggtgcaggtg-------------gtgatgag----gccccgaccccccaacccc-agGCGGGGGA-CCGT
Desmana moschata*         cagggtggga-------------gccggtagtgagacgccccgttcccagcccc-agGCGGGGGACCCTT
Galemys pyrenaicus*       ggagccggta-------------gtgacgag------gccccgttcccagcccc-agGCGGGGGACCCTT
Urotrichus talpoides*     ggtgccggtg-------------gggacgag----gcccgggtccctcaacccc-agGCGGGCGGCCCCT
Condylura cristata*       tgtgacgggg----------------------------ccccttcaccaacccc-ggGCCGGG-ACCCGC
Talpa occidentalis*       ggtgccggtg-------------gtgatgag----gccgcg---------------------------CT

Homo sapiens              GCCC--------TGCACCAGCCCCAATCAGACGCCGCTCCTCC---AACTACCGCGCTTATGCCACGGAG
Mus musculus              AGCC--------TGCGCCTGCTCCTGTCCGACGCCGCTCCTCTGCCAACTACCGAGCCTATGCCACCGAG
Solenodon paradoxus       GCCC--------AGCTCCCGCCCCTGTGCGACGCCGCTCCTCTGCCAACTACCGTGCCTACGCCACCGAA
Erinaceus europaeus       GGCC--------CGCGGCCAACCCGGTCCGACGGCGCTCCTCCGCCAACTACCGCGCCTACGCCAAGGAG
Neohylomys hainanensis    GCCC--------CCCACCCAACCCGGTCCGACGGCGCTCCTCTGCCAACTACCGCGCCTACGCCACGGAA
Uropsilus gracilis*       GCCC--------CGTGCCAGCTCCGTCCCACCTGATC---TCGGCCATCTACCCTGTCTATGCCAAGGAG
Uropsilus nivatus*        GCCC--------CGTGCCAGCTCCGTCCCACCTGATC---TCGGCCATCTACCCTGTCTATGCCAAGGAG
Scalopus aquaticus*       GCCC--------CGCGCCGGCTTTGGTGCGACGCCGCTCCCCGGCCAACTACCGCGCCTGCGCCACGG-G
Desmana moschata*         GCCT--------CGCGCCCGCTCTGGTGCGACGCCGCTCCTCTGCCAACTACCGCGCCTCCACCACGGAG
Galemys pyrenaicus*       GCCC--------CGCGCCCGCTCTGGTGCGACGCCGCTCCTCCCCCAACTACCGCGCCTCCACCACGGAG
Urotrichus talpoides*     GCCCACGCAGCGGGCTCTGGCTCTGGCGCGACGCCACTCCTCCGCCCAC------------GCCACGGAG
Condylura cristata*       GCCC--------CGCGCCGGCTCTGATGCGACCCCACCTCTCCGCCAAGTACCGCGCCTCCACCACGGAG
Talpa occidentalis*       CCCC--------CGCGCCGGCTCCGGTGCG----------TCGGCCGACCACCGCGCC-GCGCCACGGAG

Homo sapiens              CCGCACGCCAAGgtg-ggacggggcttcctgggggcagagtacaggcgccggaggg-atc-caagaccct
Mus musculus              CCACACGCCAAGgtg-gggcggggtccaagaggatgggcttctgggttcaagaatc-ctc-tgagaccca
Solenodon paradoxus       CCGCATGCCAAGgtgagggcgggg-ccagcgcggtgggcggggcctgacagcctag-agttggccatcac
Erinaceus europaeus       CCGCACGCCAAGgtgagggcgggg-cctgcatgcagataaggccacgcgccgggagccgtcctcctcc--
Neohylomys hainanensis    CCCCACGCCAAGgtgagggcgagg-cctgcgtgcccactaggtcacgcccctgggcgagagcggggccac
Uropsilus gracilis*       CCTCACGCCARGgtg-gggcgcgg-cctcccggct--------cgggatgcagatt-ctc--aggacggt
Uropsilus nivatus*        CCTCACGCCAAGgtg-gggcgcgg-cctcccggct--------cgggatgcagatt-ctc--aggayggt
Scalopus aquaticus*       GCTCGCGCCAAGgtgagggcgcgg-ccagcccgagggcgcaaccgggacgcggc---ctc-cagggtcgg
Desmana moschata*         CCTCACGCCAAGgtgagggcgcgg-ccagccggagggcgccact-cgacgcggc---ctc-cagggtcca
Galemys pyrenaicus*       CCTCACGCCAAGgtgagggcgcgg-ccagccggagggcgccacc-cgacgcggc---ctg-cagggtcca
Urotrichus talpoides*     CCTCAAGCCGCGgtgagggtgcgg-cccgcccgtgggagtcgg--attcgcagt------ggggggaccc
Condylura cristata*       CCTCACGCCAGGgtgagggcgcgg-cgggcccgagggcgcgacccggacgcggc---ctc-tagggtgga
Talpa occidentalis*       CCGCTCGCCAAGgtcagaggcggc-cgggcgcccggggggcggggtttcgaggc---tggggcgggaccg
```

**Figure S4. Genomic DNA alignment illustrating the different mechanisms of *TNNI3* exon 3 inactivation in the talpid mole lineage (mole species are denoted by asterisks; the two desman species with an intact exon 3 are underlined).** The coding region of exon 3 is provided in uppercase letters, while the 5' and 3' flanking intron sequences are provided in lowercase letters. The acceptor ('ag') and donor ('gt') splice sites are shown in bold, with mutations in red font. The codons encoding the PKA phosphorylation site ($Ser_{23,24}$) in species with an intact exon 3 are presented in underlined italics. Frameshift deletions are highlighted in pink; a single frameshift insertion is highlighted in cyan; and a deletion that encompasses the intron 3-exon 3 boundary in *Talpa occidentalis* is highlighted in green. The *T. occidentalis* intron 3-exon 3 and upstream 10 bp deletions are also evident in a single read from the low coverage *T. europaea* SRA (SRR11679552) file (data S1).

A

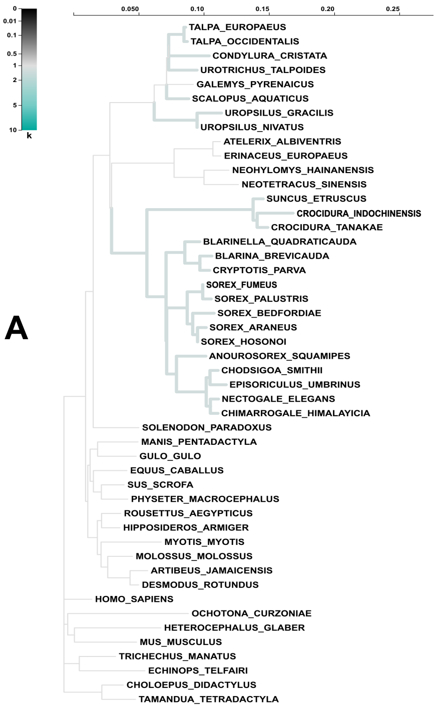

B

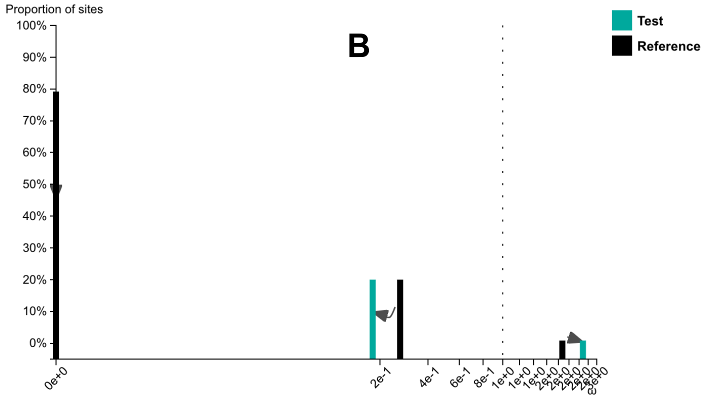

Test for selection **intensification** (K = **1.32**) was **not significant** (p = **0.118**, LR = **2.44**).

C

**Model fits**

| Model | *log* L | #. params | $AIC_c$ | Branch set | $\omega_1$ | $\omega_2$ | $\omega_3$ |
|---|---|---|---|---|---|---|---|
| RELAX alternative | -5616.8 | 113 | 11462.5 | Test | 0.00 (79.17%) | 0.18 (20.00%) | 2.28 (0.83%) |
| | | | | Reference | 0.00 (79.17%) | 0.27 (20.00%) | 1.87 (0.83%) |
| RELAX null | -5618.1 | 112 | 11462.9 | Test | 0.01 (91.69%) | 0.65 (8.31%) | 1.14 (0.00%) |
| | | | | Reference | 0.01 (91.69%) | 0.65 (8.31%) | 1.14 (0.00%) |

**Figure S5. RELAX (55) results for the analysis with *TNNI3* coding exons 1, 2, and 4-8 with 48 placental mammals.** The 'Test' or foreground category includes stem and crown soricid (shrew) branches and all stem and crown branches within Talpidae (moles) that have inactivating mutations (also see data S2). All other branches belong to the 'Reference' or background category. (A) Phylogenetic tree showing Test branches (thick lines) versus Reference branches (thin lines). The scale bar for the phylogenetic tree is in substitutions/site. The color of the Test branches corresponds to the scale bar for k in the upper left corner, where k is the selection intensity parameter. Further, k is the power to which ω for the Reference branches ($\omega_R$) is raised so that it will be equal to ω for the Test branches ($\omega_T$), i.e., $\omega_T = \omega_R$. Selection on the Test branches can either be intensified (k > 1) or relaxed (k < 1) relative to the Reference branches, and the RELAX program uses a lnL ratio test to test for significance. (B) Omega plot showing ω values and frequencies for three classes of ω: ω1, ω2, and ω3. (C) Model fits for the NULL and RELAX models and results of the lnL test. The test for differences in selection between the two branch categories was not significant (p = 0.118).

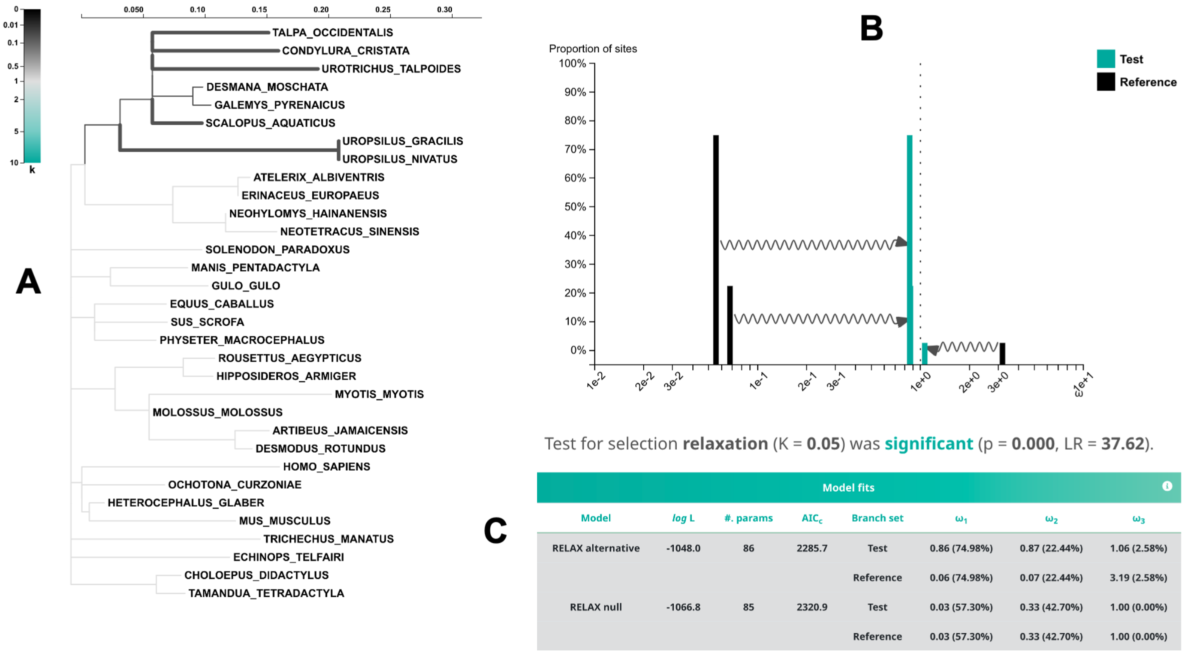

| Model | log L | #. params | AIC$_c$ | Branch set | ω$_1$ | ω$_2$ | ω$_3$ |
|---|---|---|---|---|---|---|---|
| RELAX alternative | -1048.0 | 86 | 2285.7 | Test | 0.86 (74.98%) | 0.87 (22.44%) | 1.06 (2.58%) |
| | | | | Reference | 0.06 (74.98%) | 0.07 (22.44%) | 3.19 (2.58%) |
| RELAX null | -1066.8 | 85 | 2320.9 | Test | 0.03 (57.30%) | 0.33 (42.70%) | 1.00 (0.00%) |
| | | | | Reference | 0.03 (57.30%) | 0.33 (42.70%) | 1.00 (0.00%) |

**Figure S6. RELAX (55) results for the analysis with *TNNI3* coding exons 3 with 32 placental mammals.** Sixteen shrews were not included in this analysis because any remnants of exon 3 are not recognizable. Inactivating frameshifts in talpid pseudoexons were filled in with Ns to restore complete codons. The 'Test' or foreground category includes all stem and crown branches within Talpidae (moles) that have inactivating mutations (also see Data S2). The 'Reference' or background category includes all non-talpid branches. This analysis also included Unclassified branches, which were excluded from the ω calculations and statistical comparison (also see data S2). (A) Phylogenetic tree showing Test branches (thick lines), Reference branches (thin, light lines), and Unclassified branches (thin, darker lines). The scale bar for the phylogenetic tree is in substitutions/site. The color of the Test branches corresponds to the scale bar for k in the upper left corner, where k is the selection intensity parameter. Further, k is the power to which ω for the Reference branches ($\omega_R$) is raised so that it will be equal to ω for the Test branches ($\omega_T$). Selection on the Test branches can either be intensified ($k > 1$) or relaxed ($k < 1$) relative to the Reference branches, and the RELAX program uses a lnL ratio test to test for significance. (B) Omega plot showing ω values and frequencies for three classes of ω: ω1, ω2, and ω3. (C) Model fits for the NULL and RELAX models and results of the lnL test. The test for differences in selection between the two branch categories was significant (p = 0.000).

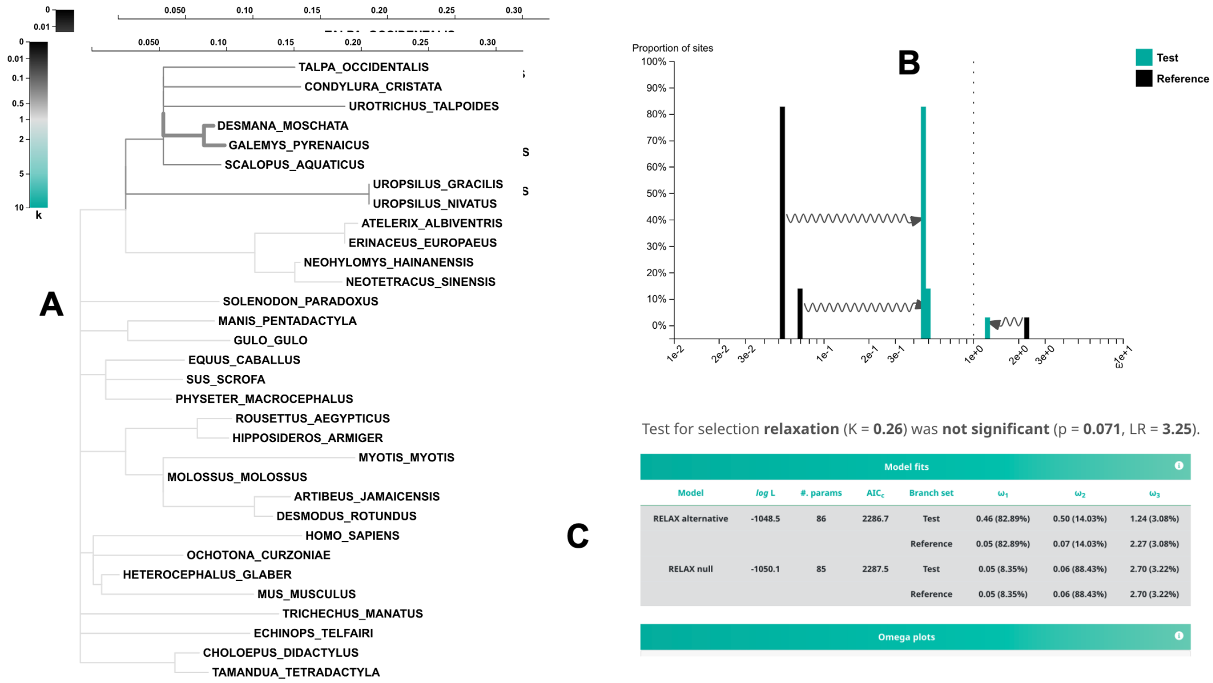

**Figure S7. RELAX (55) results for the analysis with *TNNI3* coding exons 3 with 32 placental mammals.** Sixteen shrews were not included in this analysis because any remnants of exon 3 are not recognizable. Inactivating frameshifts in talpid pseudoexons were filled in with Ns to restore complete codons. The 'Test' or foreground category includes both desmans (*Desmana*, *Galemys*) and their stem branch. The ‘Reference’ or background category includes all non-talpid branches. This analysis also included Unclassified branches, which were excluded from the ω calculations and statistical comparison (also see data S2). (A) Phylogenetic tree showing Test branches (thick lines), Reference branches (thin, light lines), and Unclassified branches (thin, darker lines). The scale bar for the phylogenetic tree is in substitutions/site. The color of the Test branches corresponds to the scale bar for k in the upper left corner, where k is the selection intensity parameter. Further, k is the power to which ω for the Reference branches ($\omega_R$) is raised so that it will be equal to ω for the Test branches ($\omega_T$). Selection on the Test branches can either be intensified ($k > 1$) or relaxed ($k < 1$) relative to the Reference branches, and the RELAX program uses a lnL ratio test to test for significance. (B) Omega plot showing ω values and frequencies for three classes of ω: ω1, ω2, and ω3. (C) Model fits for the NULL and RELAX models and results of the lnL test. The test for differences in selection between the two branch categories was not significant ($p = 0.071$).

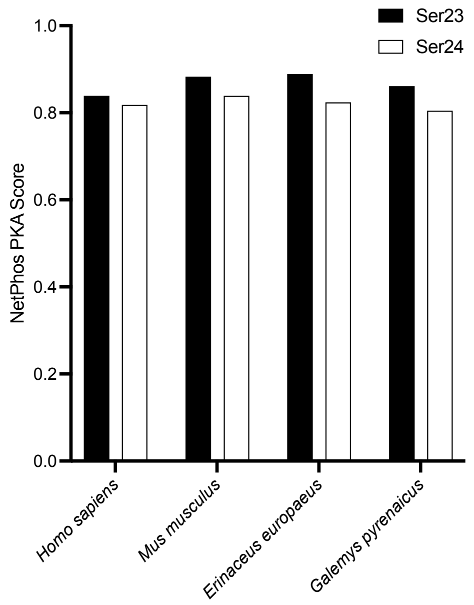

**Figure S8. Protein kinase A (PKA) target scores for Ser$_{23}$ and Ser$_{24}$ (or equivalents) in human (*Homo sapiens*), mouse (*Mus musculus*), European hedgehog (*Erinaceus europaeus*) and Pyrenean desman (*Galemys pyrenaicus*) calculated with NetPhos 3.1** (*44*). Scores show if a given amino acid residue is predicted to be a target for a specific kinase, wherein values above 0.5 are considered 'positive' hits.

**A**

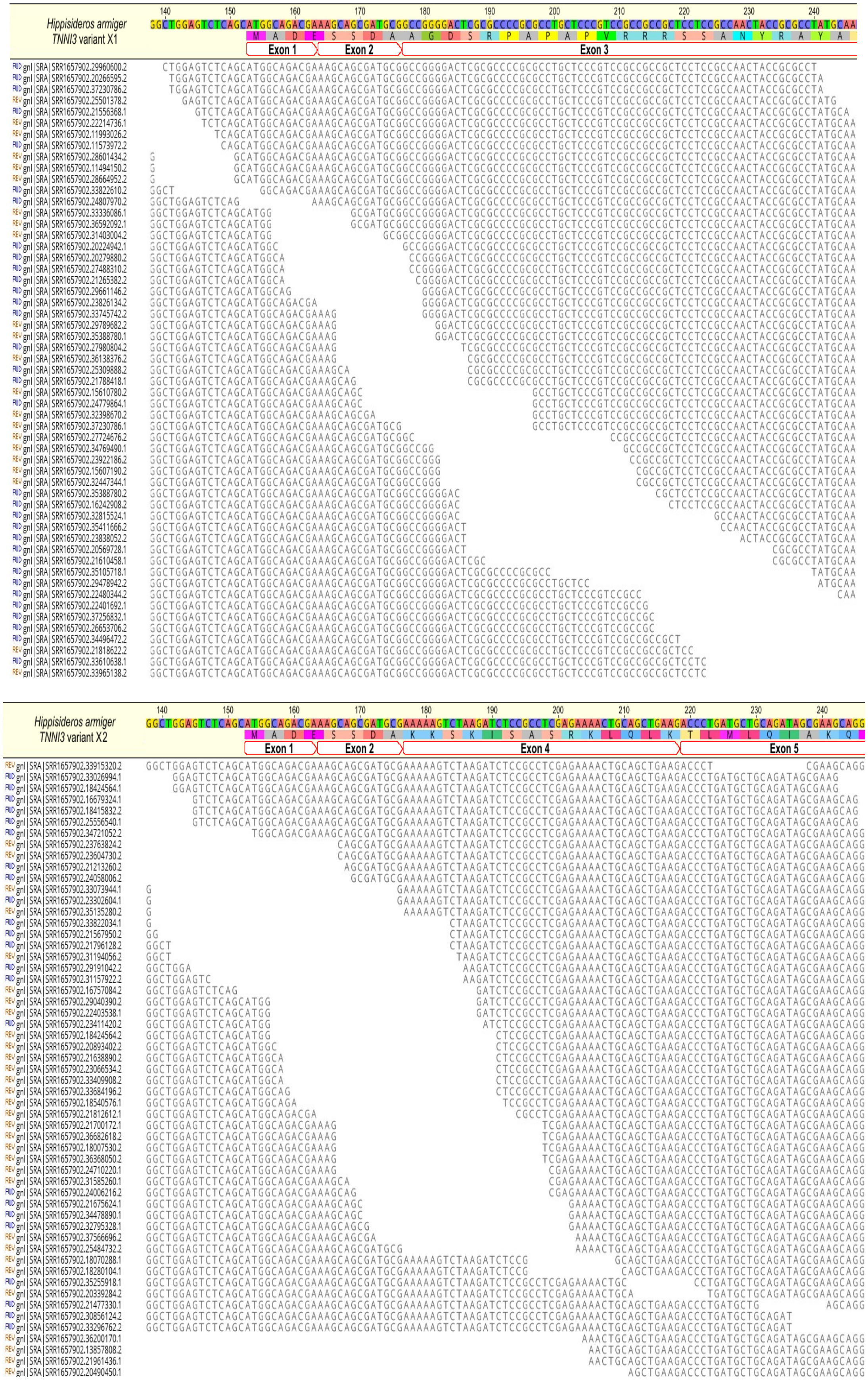

Hippsideros armiger
TNNI3 variant X1
Exon 1
Exon 2
Exon 3
Hippsideros armiger
TNNI3 variant X2
Exon 1
Exon 2
Exon 4
Exon 5

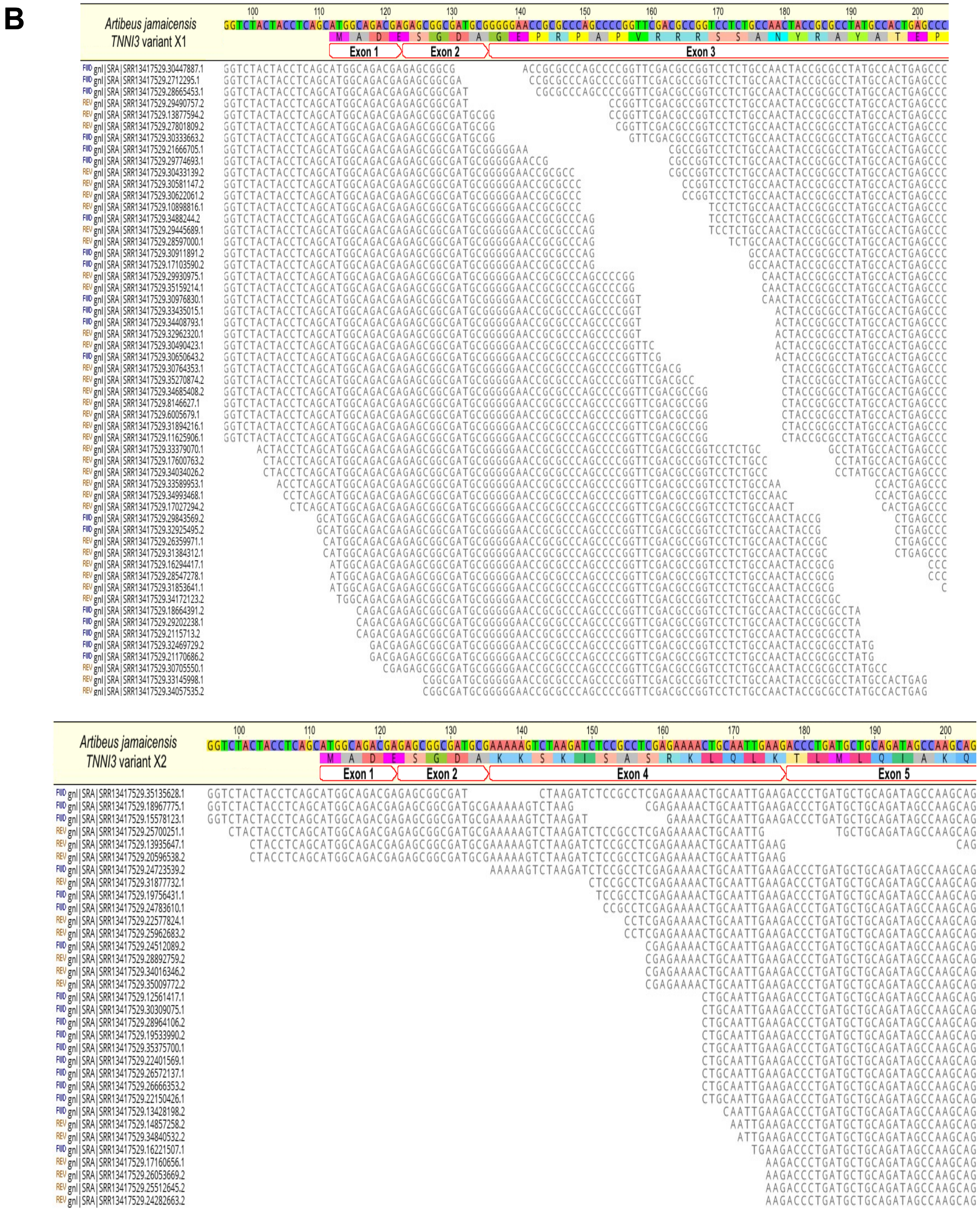

**Figure S9.** ***TNNI3*** **heart transcriptome assemblies of (A)** ***Hippisideros armiger*** **and (B)** ***Artibeus jamaicensis*** **illustrating the presence of two alternatively spliced variants.** Note that the X2 variant of both species lacks exon 3.

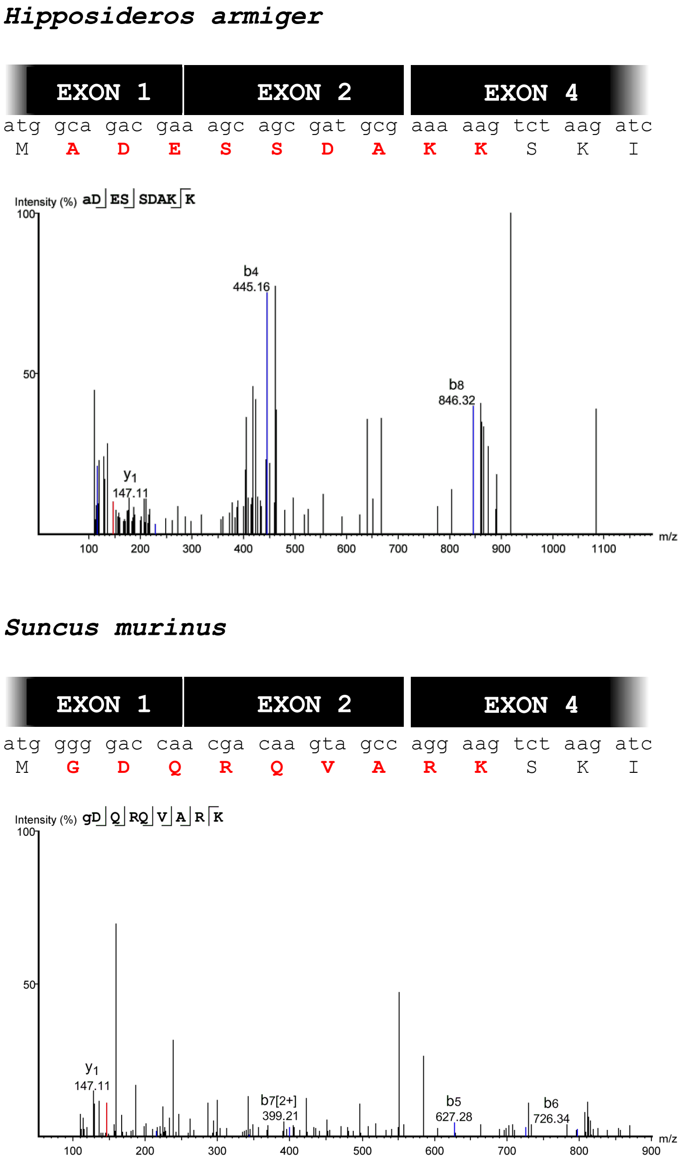

**Figure S10. Liquid chromatography with tandem mass spectrometry (LC-MS/MS) protein sequencing with parallel reaction monitoring to identify N-terminal protein sequence in ~22 kDa cTnI bands from great roundleaf bat (*Hipposideros armiger*) and Asian house shrew (*Suncus murinus*).** In both cases, peptides were identified that span *TNNI3* exons 1, 2 and 4, confirming that exon 3$^{-}$ mRNA is translated.

```
                          Ser23&24
mouse            MADESSDAAGEPQPAPAPVRRRSSANYRAYATEPHAKKKSKISASRKLQLKTLMLQIAKQ
mouse_truncated  ---------------------------MAYATEPHAKKKSKISASRKLQLKTLMLQIAKQ
common shrew     -----------------------------MADERSVARKSKITASRKLQLKTLMLQIAKQ
Iberian mole     -----------------------------MADESREAKKSKISASRKLQLKSLMLQVAKQ
                                               * *    :****:********:****:***

mouse            EMEREAEERRGEKGRVLRTRCQPLELDGLGFEELQDLCRQLHARVDKVDEERYDVEAKVT
mouse_truncated  EMEREAEERRGEKGRVLRTRCQPLELDGLGFEELQDLCRQLHARVDKVDEERYDVEAKVT
common shrew     ELEREVEERRGEKGRALNSRCQPLELAGLGFAELQDLCRQLHAQVDKVDEERYDVEAKVT
Iberian mole     ELEREAEERRGEKGRALSARCEPLELAGLGVAELQDLCRQLHARVDKVDEERYDVEAKVT
                 *:***.*********.* :**:**** ***. ***********:****************

mouse            KNITEIADLTQKIYDLRGKFKRPTLRRVRISADAMMQALLGTRAKESLDLRAHLKQVKKE
mouse_truncated  KNITEIADLTQKIYDLRGKFKRPTLRRVRISADAMMQALLGTRAKESLDLRAHLKQVKKE
common shrew     KNISEIADLTQKVIDLRGKFKRPTLRRVRISADAMMQALLGTRAKESLDLRANLKQVKKE
Iberian mole     KNVSEIADLTQKILDLRGKFKRPTLRRVRISADAMMQALLGTRAKESLDLRANLKQVKKE
                 **::********: **************************************:*******

mouse            DIEKENREVGDWRKNIDALSGMEGRKKKFEG----
mouse_truncated  DIEKENREVGDWRKNIDALSGMEGRKKKFEG----
common shrew     DTEKENREVGDWRKNIDALSGMEGRKKKFEG----
Iberian mole     DTEKENREVGDWRKNIDALSGMEGRKKKFEGSGQA
                 * *****************************
```

**Figure S11. The genetic excision of exon 3 in *TNNI3* of shrews and moles recapitulates the truncated cardiac troponin I previously generated in transgenic mouse studies**. The truncated mouse cTnI sequence is derived from Barbato et al. (2005) (*22*) who created a translation initiation codon before $Ala_{29}$. Shrews are represented by the common shrew, *Sorex araneus*, and moles are represented by the Iberian mole, *Talpa occidentalis* (see data S1 for sequence accession information). Red arrows denote the pertinent protein kinase A (PKA) target residues ($Ser_{23/24}$).
